# Targeting the glycan epitope type I *N*-acetyllactosamine enables immunodepletion of human pluripotent stem cells from early differentiated cells

**DOI:** 10.1101/2023.08.30.555456

**Authors:** Charlotte Rossdam, Smilla Brand, Julia Beimdiek, Astrid Oberbeck, Marco Darius Albers, Ortwin Naujok, Falk F. R. Buettner

## Abstract

Cell surface biomarkers are fundamental for specific characterization of human pluripotent stem cells (hPSCs). Importantly, they can be applied for hPSC enrichment and/or purification but also to remove potentially teratoma-forming hPSCs from differentiated populations before clinical application. Several specific markers for hPSCs are glycoconjugates comprising the glycosphingolipid (GSL)-based glycans SSEA-3 and SSEA-4. We applied an analytical approach based on multiplexed capillary gel electrophoresis coupled to laser-induced fluorescence detection to quantitatively assess the GSL glycome of human embryonic stem cells and human induced pluripotent stem cells as well as during early stages of differentiation into mesoderm, endoderm, and ectoderm. Thereby, we identified the GSL lacto-*N*-tetraosylceramide (Lc4-Cer, Galβ1-3GlcNAcβ1-3Galβ1-4Glc-Cer), which comprises a terminal type 1 LacNAc (T1LN) structure (Galβ1-3GlcNAc), to be rapidly decreased upon onset of differentiation. Using a specific antibody, we could confirm a decline of T1LN-terminating glycans during the first four days of differentiation by live-cell staining and subsequent flow cytometry. We could further separate T1LN-positive and T1LN-negative cells out of a mixed population of pluripotent and differentiated cells by magnetic activated cell sorting. Notably, not only the T1LN-positive but also the T1LN-negative population was positive for SSEA-3, SSEA-4, and SSEA-5 while expression of nuclear pluripotency markers *OCT4* and *NANOG* was highly reduced in the T1LN-negative population, exclusively. Our findings suggest T1LN as a pluripotent stem cell-specific glycan epitope that is more rapidly down-regulated upon differentiation than SSEA-3, SSEA-4, and SSEA-5.

## Introduction

Human pluripotent stem cells (hPSCs) including both, human embryonic stem cells (hESCs) as well as human induced pluripotent stem cells (hiPSCs), offer great potential in regenerative medicine as an unlimited cell source that can be differentiated into nearly every somatic cell type (Okano and Yamanaka, 2014; Pera and Trounson, 2004). hPSC-specific cell surface markers are highly relevant for example to define pluripotency and purity of cultured cells. Furthermore, pluripotency-specific markers can be applied for killing or removal of hPSCs from differentiated populations since potentially tumorigenic pluripotent cells present a major hurdle for clinical application of hPSC-derived somatic cells (Fong et al., 2010).

Numerous routinely used targets for characterization of hPSCs are cell surface exposed glycoconjugates (Wright and Andrews, 2009) such as the structures detected by the monoclonal antibodies Tra-1-60 and Tra-1-81 that have been shown to recognize a tetrasaccharide epitope which is part of a mucin-type *O*-glycan in hESCs (Natunen et al., 2011). Furthermore, glycosphingolipids (GSLs) belonging to the globo-series (Gb) such as globopentaosylceramide (Gb5-Cer, SSEA-3), sialyl-globopentaosylceramide (sialyl-Gb5-Cer, SSEA-4) (Kannagi et al., 1983), and fucosyl-globopentaosylceramide (fucosyl-Gb5-Cer, Globo H) (Barone et al., 2013) or to the lacto-series (Lc) such as fucosyl-lactotetraosylceramide (fucosyl-Lc4-Cer, Fuc1-2Galβ1-3GlcNAcβ1-3Galβ1-4Glc-Cer) (Liang et al., 2010; Liang et al., 2011) or sialyl-lactotetraosylceramide (Sia-Lc4-Cer, Siaα2-3Galβ1-3GlcNAcβ1-3Galβ1-4Glc-Cer) (Barone et al., 2014) were described to be highly specific for hPSCs and rapidly disappear from the cell surface upon differentiation.

Liang *et al*. described a general switch of the core structures of GSLs from globo- and lacto-to ganglio-series upon hPSC differentiation. This switch was accompanied by changes in expression levels of the involved glycosyltransferases (Liang et al., 2010; Liang et al., 2011). Both, globo- and lacto-series GSLs are characterized by a galactose bound in β1-3-linkage to either *N*-acetylgalactosamine (Galβ1-3GalNAc) or *N*-acetylglucosamine (Galβ1-3GlcNAc), respectively. The Galβ1,3GlcNAc disaccharide, which is commonly referred to as type 1 LacNAc chain (T1LN) has been found on hPSC-specific *O*-glycans, where it is part of the Tra-1-60 and Tra-1-81 epitopes (Natunen et al., 2011), as well as on *N*-glycans (Hasehira et al., 2012; Konze et al., 2017). Appearance of these structures is likely associated with the hPSC-specific expression of the major β1,3-galactosyltransferase (B3GALT5) (Tateno et al., 2011).

By screening of antibody libraries, an hPSC-specific antibody targeting the glycan structure Fuc1-2Galβ1-3GlcNAcβ, which is designated as H type-1 (H-1), was found. The H-1 epitope also contains a T1LN motif. This H-1-epitope-specific antibody, termed SSEA-5, was shown to detect protein *N*- and *O*-glycans (Tang et al., 2011). The H-1 epitope is also part of the lactoseries GSL fucosyl-Lc4-Cer whose levels considerably decline upon differentiation of hPSCs into embryoid bodies (EBs) (Liang et al., 2010). Tang *et al*., demonstrated the potential of the H-1 epitope to be immunologically targeted for prospective removal of hPSCs from mixed populations. hPSCs that were spiked into a population of hPSC-derived, fully differentiated cells could be efficiently removed by fluorescence-activated cell sorting (FACS) using the SSEA-5 antibody. This completely abolished teratoma-formation potential of the SSEA-5-immunodepleted population. However, applying the same sorting strategy for a pure population of day-3 differentiated hPSCs gave rise to an immunodepleted population which still caused teratoma formation. It was necessary to target two additional hPSC-specific cell surface proteins to finally select for cells that did not cause teratoma (Tang et al., 2011). These findings show that, until now, individual markers like H-1 are not sufficient for complete separation of hPSCs from early differentiated progeny, pointing towards the necessity for the identification of additional hPSC-specific cell surface markers.

We have established multiplexed capillary gel electrophoresis coupled to laser-induced fluorescence detection (xCGE-LIF) as a powerful technique for the analysis of GSL glycosylation (Cumin et al., 2022; Jirmo et al., 2020; Rossdam et al., 2019; Starzonek et al., 2020) and used this approach to screen for hPSC markers. A quantitative comparison of GSL glycan levels in hPSCs (hESCs and hiPSCs) and upon early lineage-specific differentiation revealed the lactoseries GSL lacto-*N*-tetraosylceramide (Lc4-Cer, Galβ1-3GlcNAcβ1-3Galβ1-4Glc) to be significantly down-regulated in hPSCs upon differentiation. Lc4-Cer contains a T1LN motif and by applying a T1LN-specific antibody, we confirmed surface expression of T1LN-containing glycans by flow cytometry. Using this antibody for magnetic activated cell sorting (MACS), we could quantitatively separate T1LN-expressing hPSCs from early differentiated cells. The comparison of T1LN glycans to other well-known hPSC surface markers, such as SSEA-3, SSEA-4 or H-1 corroborated the suitability of T1LN as a superior marker for undifferentiated hPSCs.

## Results

### xCGE-LIF-based profiling of GSL glycosylation reveals Lc4-Cer as a hPSC marker

We applied xCGE-LIF to analyze GSL glycosylation of hPSCs (hESCs and hiPSCs) and during early stages of differentiation with the aim to identify potential novel GSL-based biomarkers of hPSCs. Therefore, we differentiated both cell lines for 4 days into early stages of mesoderm, endoderm, and ectoderm and confirmed successful lineage specification on day 4 of differentiation (d4) by flow cytometric analyses of lineage-specific differentiation markers (Suppl. Figure S1). Differentiation into mesoderm resulted in the characteristic balanced cell surface expression of the adhesion molecules NCAM and EpCAM as described by Kempf *et al*. (Kempf et al., 2016b). Endodermal differentiation resulted in 87±5 % and 85±6 % CXCR4 positive hES-derived and hiPS-derived cells, respectively. For ectoderm, we observed 95±2 % and 97±1 % Pax-6-positive hES-derived and hiPS-derived cells, respectively (Suppl. Figure S1). For both cell lines and all differentiation approaches we harvested cells on day (d)0, d1, d2, d3, and d4 and analyzed GSL glycosylation by xCGE-LIF (Figure 1). This analysis revealed down-regulation of the known pluripotency-associated GSLs SSEA-3, SSEA-4, and fucosyl-Lc4-Cer (containing the terminal H-1 trisaccharide epitope Fuc1-2-Galβ1-3GlcNAcβ, detected by the SSEA-5 antibody) and/or globo H upon differentiation into the three germ layers. Notably, the GSLs fucosyl-Lc4-Cer (comprising the H-1 epitope) and globo H could not be discriminated as they comigrated in the capillary gel electrophoresis. However, using exoglycosidase digestions, we confirmed that the respective signal arises from both structures (data not shown). In addition to these rather expected findings, we identified the GSL glycan lactotetraose (Lc4) to be down-regulated during differentiation. Especially upon mesodermal and endodermal differentiation the decline of Lc4-levels was stronger than for the known marker structures. For ectodermal differentiation, we did not observe down-regulation of Lc4-levels. However, also the known markers showed a heterogenous profile depending on the analyzed cell line.

**Figure 1.**
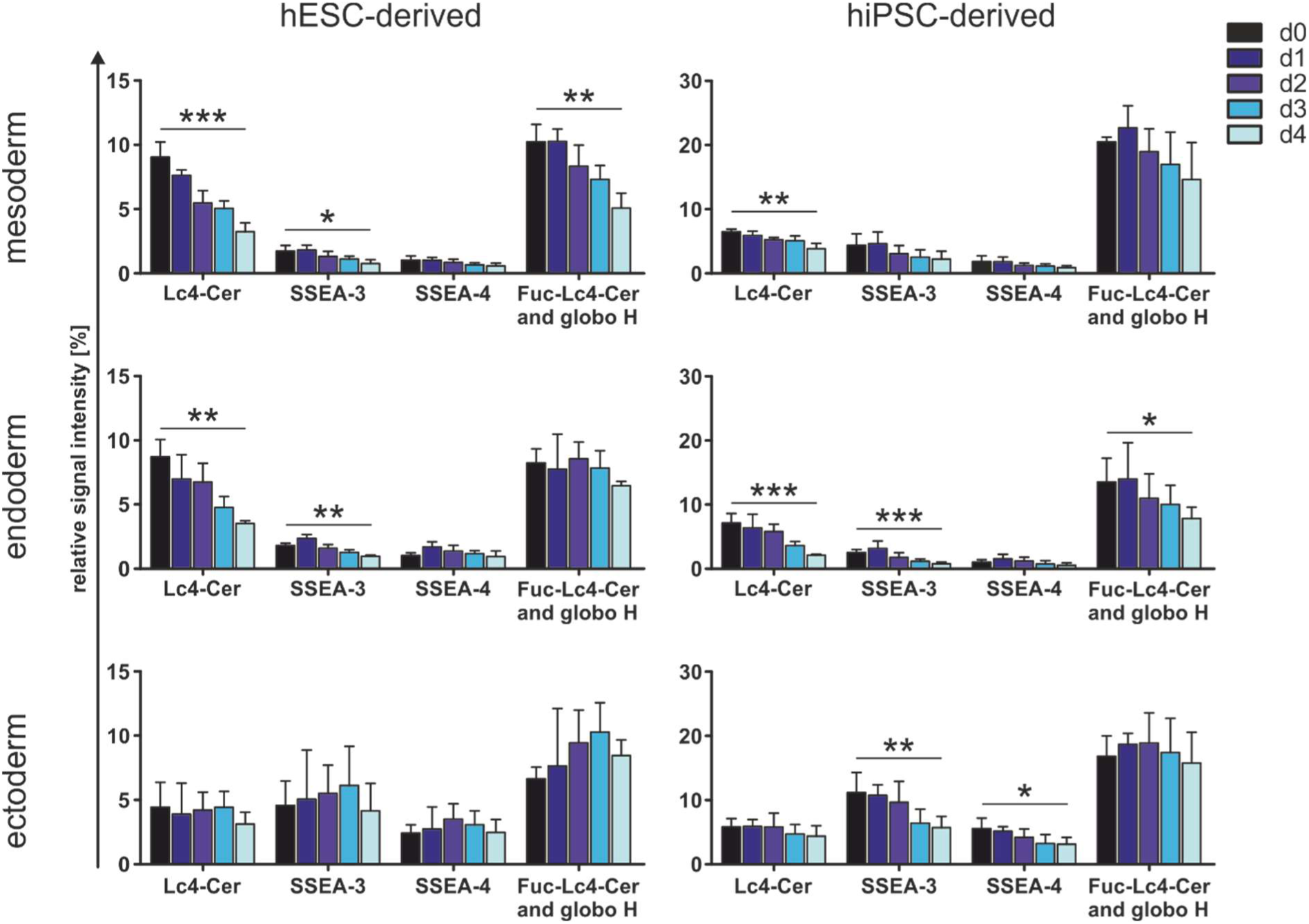
GSL analysis. xCGE-LIF analysis of APTS-labelled glycans derived from GSLs of hiPSCs (left panel) and hESCs (right panel), and their differentiation derivatives of mesoderm (top), endoderm (middle), and ectoderm (bottom) through days 0 - 4 of differentiation (d0-d4, black to light blue), respectively. Relative signal intensities are displayed for Lc4-Cer, SSEA-3, SSEA-4, and fucosyl Lc4-Cer / globo H. Bar diagrams represent mean + S.D. (*n*=3-5). An unpaired Student’s *t*-test was performed between the peaks of d0 and d4 of differentiation for each cell line and each differentiation approach. Statistically significant differences are highlighted (* *p*<0.05; ** *p*<0.01; *** *p*<0.001).

### Cell surface levels of T1LN strongly decline after four days of hPSC differentiation

As a complementary approach, we analyzed non-permeabilized hPSCs (d0) as well as their progeny on d4 of differentiation into meso-, endo-, and ectoderm by flow cytometry applying the BG-1 antibody detecting a terminal T1LN-structure (Domino et al., 2001) as present in Lc4-Cer (Figure 2, Suppl. Figure S3). The differentiation outcome was scrutinized by analysis and quantification of marker expression as described above (Suppl. Figure S2). The flow cytometric analyses revealed a strong and significant decline of T1LN cell surface levels upon differentiation for 4 days into all three lineages for both hESCs and hiPSCs (Figure 2 A, B left panels). We also assessed cell surface expression of SSEA-4 and H-1 (SSEA-5) by flow cytometry using specific antibodies. We observed a mild to moderate decrease of cell surface levels of these known markers upon differentiation. Notably, SSEA-4 and H-1 down-regulation was for all germ layers less pronounced than down-regulation of T1LN (Figure 2 A, B middle and right panels). Taken together, our findings strongly suggest T1LN as a highly specific cell surface marker for undifferentiated hPSCs that is characterized by rapid down-regulation upon differentiation.

**Figure 2.**
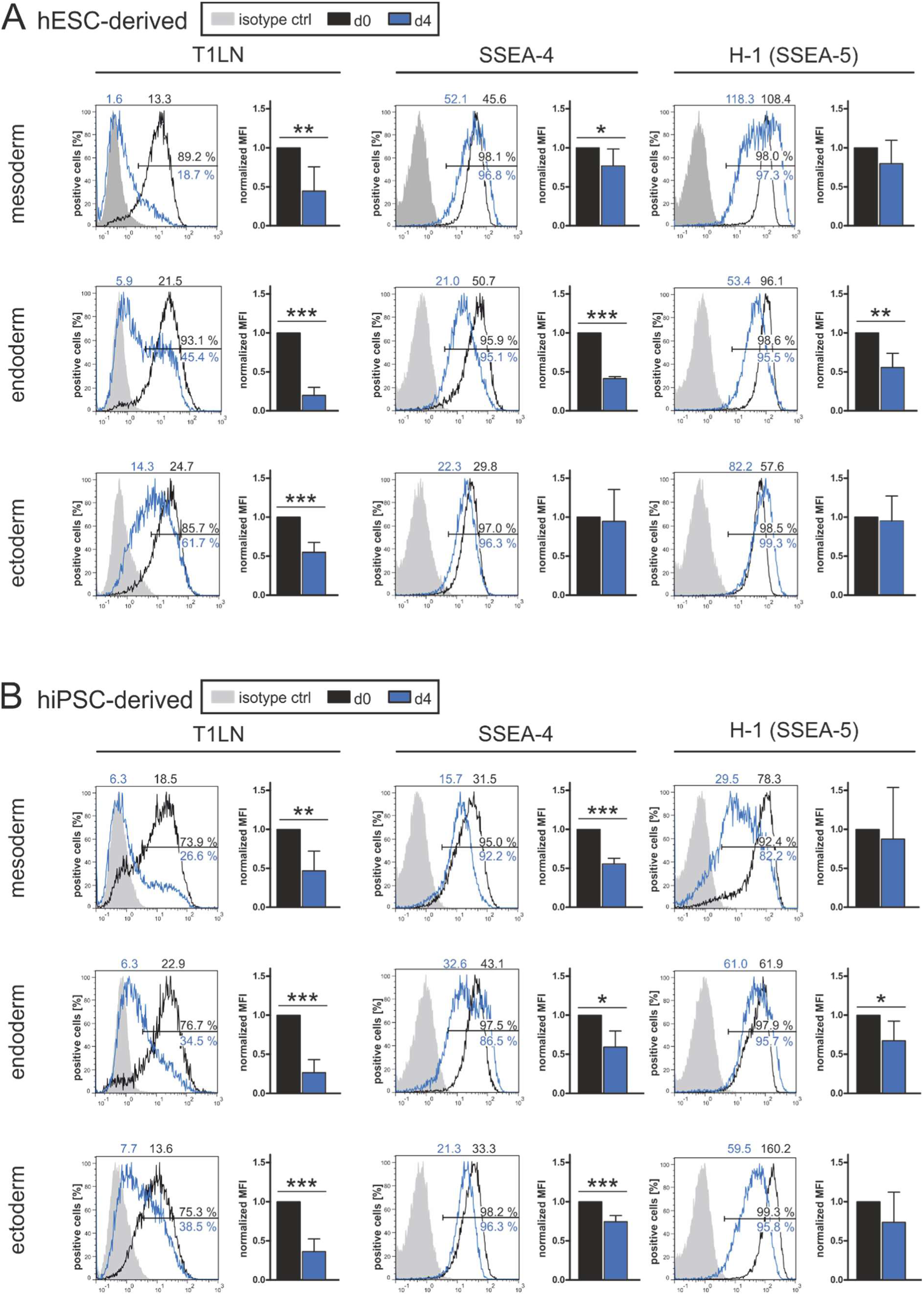
Cell surface expression of T1LN, SSEA-4, and SSEA-5. (A-B) Flow cytometry analysis of hESCs (A) and hiPSCs (B) on d0 and d4 of differentiation for each differentiation approach (top: mesoderm; middle: endoderm; bottom: ectoderm) for T1LN (left panel), SSEA-4 (middle panel), and SSEA-5 (right panel). Representative overlay histograms show fluorescence signal of viable cells on d0 (black) and d4 (blue). Cells were defined positive if the fluorescence signal was above that of 99 % of the cells stained with the respective isotype control (grey). Only isotype controls for d0 are displayed, but for calculation of positive cells on d4, isotype controls of d4 were considered. Number of positive cells are shown below (blue) and above (black) the line representing the gate. Mean fluorescence intensity is shown above histograms (d0, black and d4, blue). Bar diagrams show mean fluorescence intensities (MFI) + S.D. (*n*=3-6) normalized to d0. An unpaired Student’s *t*-test was performed for each target between d0 and d4 of differentiation and statistically significant differences are highlighted (* *p*<0.05; ** *p*<0.01; *** *p*<0.001).

### T1LN-expressing cells could be efficiently depleted by magnetic cell sorting from mixed populations

Cell surface markers enabling the immunological identification and prospective depletion of hPSCs, such as SSEA-5 described by Tang and colleagues (Tang et al., 2011), are highly demanded in regenerative medicine, since residual contaminating hPSCs within differentiation cultures might cause the formation of teratoma or teratocarcinoma upon transplantation into human recipients. With the aim to scrutinize whether the T1LN glycan epitope is useful in the above described setting, we again differentiated hPSCs (hESCs and hiPSCs) for 4 days into early meso-, endo-, and ectoderm and confirmed successful differentiation for both cell lines into all germ layer differentiations (Suppl. Figure S4). We repeated the flow cytometric analyses of d0 and d4 cells for T1LN, SSEA-4, H-1 (SSEA-5), and additionally analyzed SSEA-3 surface expression (Figure 3B). This analysis basically mirrored our previous findings (Figure 2) that the expression of T1LN as well as of known hPSC surface markers was significantly reduced on d4 of differentiation (Figure 3B, Suppl. Figure S5). To further evaluate whether hPSCs could be depleted from mixed populations based on their T1LN expression, we pooled equal cell numbers of hPSCs (hESCs or hiPSCs, d0) and their mesodermal, endodermal or ectodermal derivatives (d4, Figure 3A). Flow cytometric analyses of surface markers of the pooled population (d0/d4-pool) mostly revealed, as expected, intermediate values for fluorescence intensities (Figure 3B, Suppl. Figure S5). Using magnetic microbeads decorated with the T1LN-specific BG-1 antibody for MACS analyses, we could obtain an unbound fraction with less than 1% of cells being detected as T1LN-positive by subsequent flow cytometry (Figure 3B, Suppl. Figure S5). This fraction was designated as T1LN-negative and confirmed the successful depletion of T1LN-positive cells from the mixed population. Notably, expression of the stem cell markers SSEA-4 and H-1 (SSEA-5), were - if at all - only slightly reduced in the unbound T1LN-negative fraction in comparison to the d0/d4-pool. On the other hand, levels of SSEA-3 and T1LN were significantly diminished (Figure 3B, Suppl. Figure S6). All analysed surface markers were detected at high levels upon MACS in the bound fraction, which we designated as T1LN-positive. Taken together, our analyses revealed that upon germ layer differentiation a substantial fraction of cells already had lost T1LN cell surface expression while still carrying the pluripotency markers SSEA-3, SSEA-4, and H-1 (SSEA-5) on their cell surface (Suppl. Figure S6).

**Figure 3.**
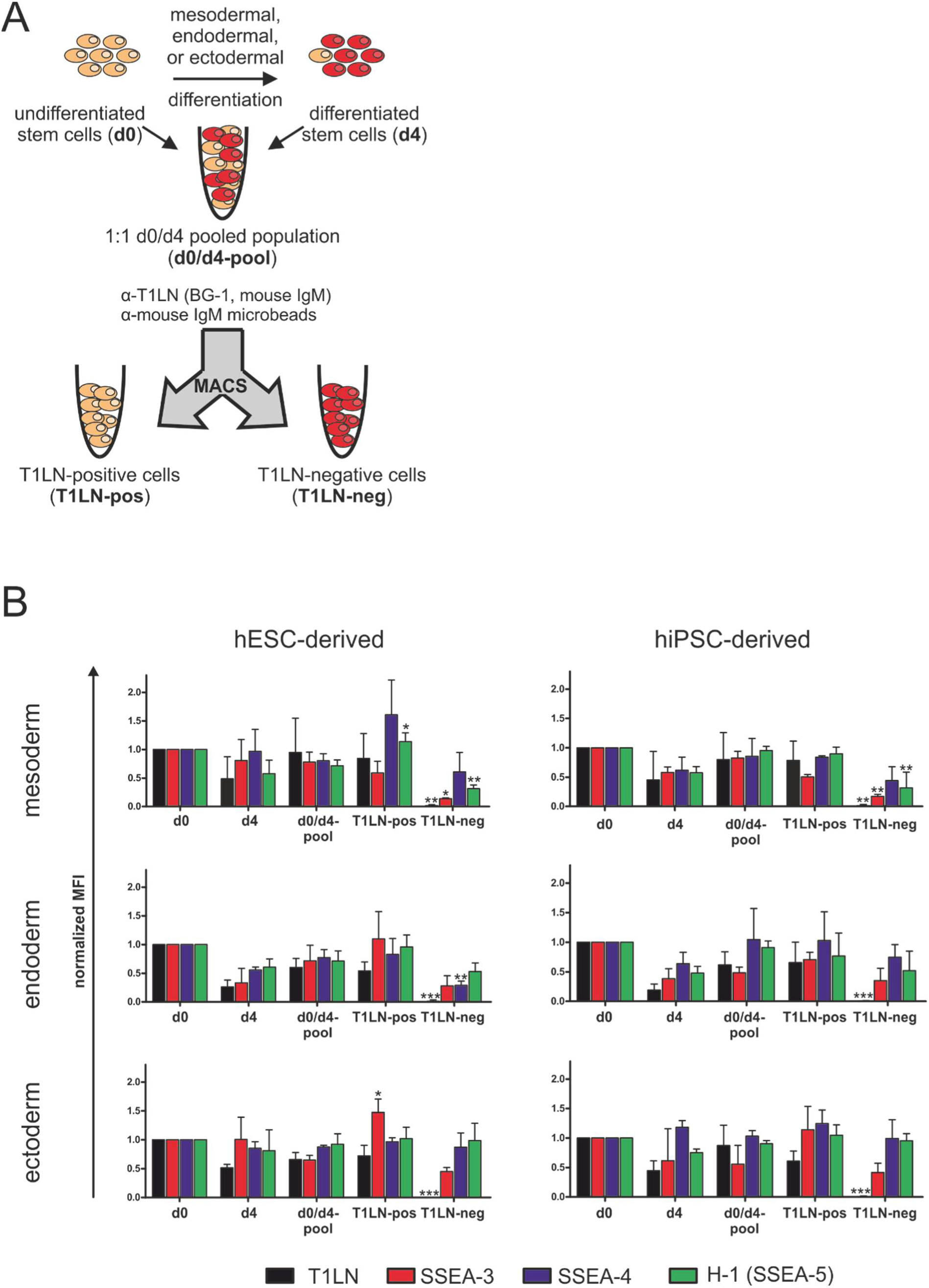
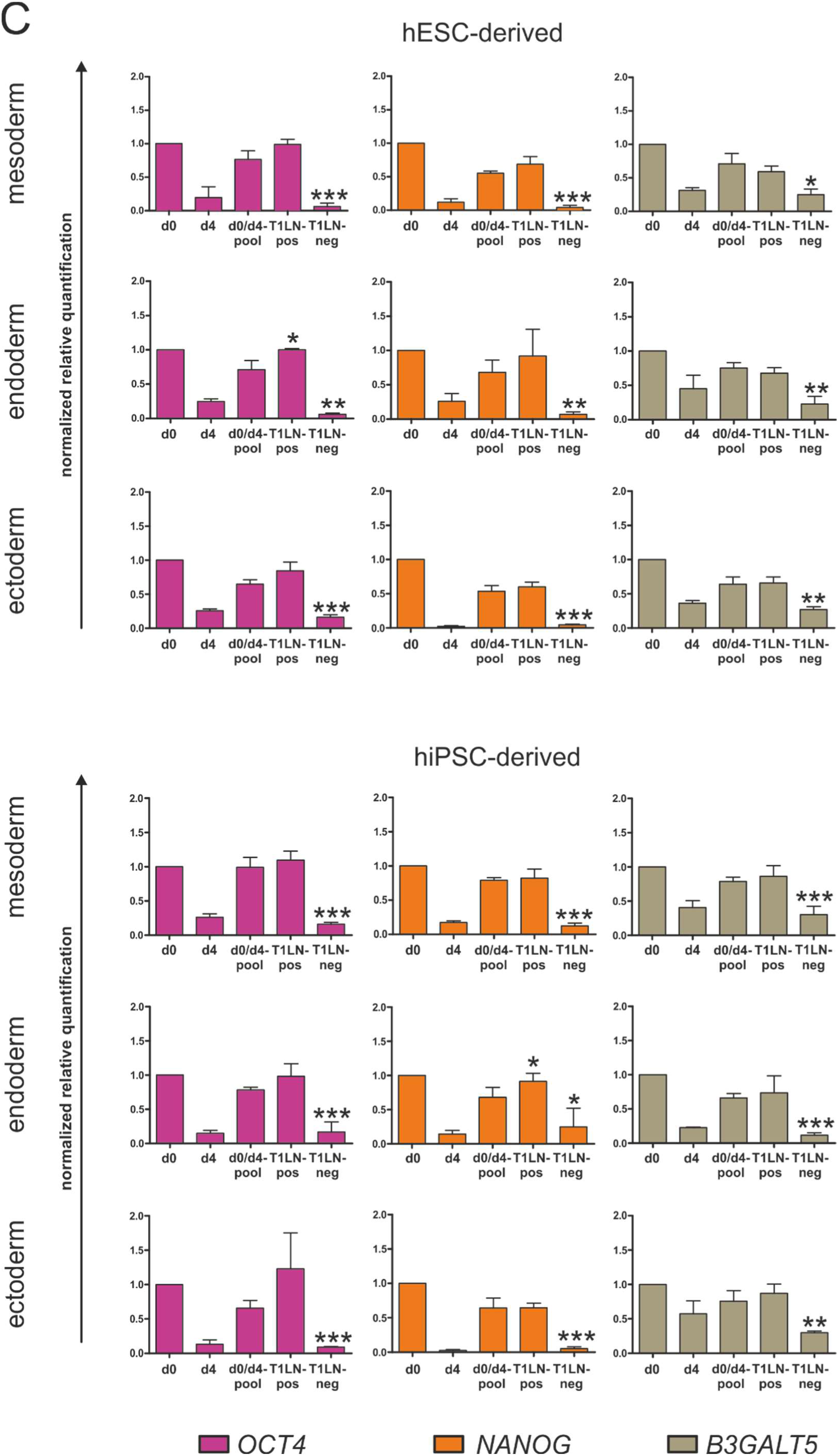
Cell sorting and analysis of T1LN, SSEA-3, SSEA-4, and SSEA-5. (A) Schematic illustration of MACS assay using a 1:1 pool of undifferentiated stem cells (d0, shown in orange) and differentiated stem cells (d4, shown in red) (d0/d4-pool). Cell sorting using the mouse IgM antibody BG-1 directed against T1LN and α-mouse IgM microbeads generated a pool of T1LN-positive cells (T1LN-pos) and T1LN-negative cells (T1LN-neg). (B) Cell surface expression of T1LN, SSEA-3, SSEA-4, and SSEA-5 in hPSCs and differentiated derivatives pre- and post-MACS using the α-T1LN antibody. Bar diagrams show mean fluorescence intensities (MFI, normalized to d0) + S.D. (*n*=3-7) of T1LN (black), SSEA-3 (red), SSEA-4 (blue), and SSEA-5 (green). (C) Relative expression of *OCT4*, *NANOG,* and β-1-3-galactosyltransferase *B3GALT5* in hPSCs and differentiated derivatives pre- and post-MACS analyzed by qPCR. Error bars represent mean + S.D. (*n*=3-4). For both B and C, expression was analyzed in hESCs (left panel) and hiPSCs (right panel) and their differentiated derivatives (top: mesoderm; middle: endoderm; bottom: ectoderm) pre-MACS in d0, d4 and d0/d4-pooled cells, as well as post-MACS in the T1LN-positive and T1LN-negative population. An unpaired Student’s *t*-test was performed for each target between d0/d4-pool and T1LN-pos or T1LN-neg, respectively, and statistically significant differences are highlighted (* *p*<0.05; ** *p*<0.01; *** *p*<0.001).

### Expression of *OCT4* and *NANOG* is strongly reduced in the T1LN-negative population

The transcription factors OCT4 and NANOG are generally considered as gold standards for pluripotency of hPSCs. We therefore assessed their gene expression by quantitative PCR (qPCR) in the different populations comprising hPSCs (d0), differentiated cells (d4), pooled cells (d0/d4-pool), and upon MACS of the sorted cells (T1LN-pos and T1LN-neg). This analysis expectedly revealed high expression of *OCT4* and *NANOG* in d0 hPSCs which was drastically diminished upon 4 days of differentiation (d4) for both cell lines and all differentiation approaches (Figure 3C). The d0/d4-pool showed an expected intermediate expression of *OCT4* and *NANOG*. Upon cell sorting by MACS, we observed high expression of these markers in the T1LN-positive population and a strongly reduced expression in the T1LN-negative population which was even lower than on d4. This finding suggests that the T1LN-negative population is highly depleted of pluripotent cells. As we previously showed that the T1LN-negative population is still positive for SSEA-3, SSEA-4, and H-1 (SSEA-5), T1LN seems to be a more sensitive cell surface pluripotency maker than the other commonly applied glycan markers.

### T1LN cell surface levels are correlated to *B3GALT5* gene expression

The cell surface glycan epitopes T1LN, SSEA-3, SSEA-4, and H-1 (SSEA-5) all contain a galactose bound in β1-3-linkage to either GlcNAc or GalNAc which is synthesized by the enzyme B3GALT5 (Zhou et al., 2000). Thus, we also determined *B3GALT5* expression by qPCR in our pre- and post-MACS cell populations (d0, d4, d0/d4-pool, T1LN-pos, T1LN-neg). Thereby a decrease of *B3GALT5* mRNA expression on d4 vs. d0 for both cell lines and mesodermal, endodermal, and ectodermal differentiations was observed. *B3GALT5* expression appeared to be even more strongly reduced in the post-MACS T1LN-neg cell population than on d4 of differentiation hinting towards a strong correlation of T1LN levels and *B3GALT5* expression. (Figure 3C).

## Discussion

Ideal surface markers for hPSCs correlate with the pluripotent state and can be efferently targeted by specific antibodies. Thereby such markers can be applied to examine pluripotency but also for antibody-based enrichment or depletion of hPSCs. In particular, immunodepletion of hPSCs from *in vitro* differentiated populations has gained attention as residual hPSCs might give rise to teratoma formation upon clinical application in human (Fong et al., 2010). Strategies for depletion of hPSCs for example comprise specific cytotoxic antibodies (Choo et al., 2008; Tan et al., 2009), prospective removal by fluorescence-activated cell sorting (Tang et al., 2011) and suicide gene-mediated ablation of pluripotent cells (Naujok et al., 2010). To prevent teratoma formation in mice, a combination of different cell surface markers was necessary for comprehensive immunodepletion of hPSCs from early differentiated populations (Tang et al., 2011), highlighting the need to search for further highly specific hPSC markers.

Many widely applied cell surface markers for hPSC characterization are glycoconjugates such as the *O*-glycan epitopes Tra-1-60 and Tra-1-81 (Natunen et al., 2011) or the globoseries GSLs SSEA-3 and SSEA-4 which were long known cell surface markers of stem cells and teratocarcinoma cells (Carpenter et al., 2004; Kannagi et al., 1983). Furthermore, the glycan epitope sialyl-Lc4 was shown to be present on glycoproteins and GSLs of hPSCs and this glycan structure rapidly decreased upon differentiation into hepatocyte- and cardiomyocyte-like cells (Barone et al., 2014). Additionally, the H-1 epitope which can build the terminal structure of *O*-glycans, *N*-glycans and GSL-glycans (e.g. Fuc-Lc4-Cer) was discovered by Liang *et al*. to be down-regulated upon hPSC differentiation (Liang et al., 2010). Shortly after, Tang *et al*. introduced the hPSC-specific anti-SSEA-5 antibody directed against the H-1 glycan epitope and well suited for immunodepletion of hPSCs from mixed cell populations (Tang et al., 2011).

GSLs comprise a highly diverse family of glycosylated membrane constituents whose synthesis is tightly controlled by the cellular state and during cell fate specification, rendering them attractive marker candidates (D’Angelo et al., 2013; Russo et al., 2018). When we recently profiled changes in GSL glycosylation upon cardiomyogenic differentiation of hPSCs, we not only noted strongly reduced levels of the known markers SSEA-3, SSEA-4, globo H, and Fuc-Lc4-Cer (comprising the H-1 epitope which can be detected by the anti-SSEA5 antibody), but also of its biosynthetic precursor Lc4-Cer (Rossdam et al., 2019). Therefore, we assessed GSL glycosylation of hPSCs and during early stages of lineage-specific differentiation applying our novel xCGE-LIF-based analytical approach. This analysis revealed that Lc4-Cer levels rapidly decline upon differentiation. In 2010, Liang *et al*. first described Lc4-Cer to be down-regulated upon hESC differentiation into embryoid bodies. However, in latter and subsequent studies, Lc4-Cer could not be clearly distinguished from its structural isomers Gb4-Cer (GalNAcβ1-3Galα1-4Galβ1-4Glc-Cer) and nLc4 (Galβ1-4GlcNAcβ1-3Galβ1-4Glc) by using MALDI-MS, MALDI-MS/MS or LC-MS/MS for glycosphingolipid profiling (Fujitani et al., 2013; Ho et al., 2017; Liang et al., 2010; Liang et al., 2011; Lin et al., 2020; Ojima et al., 2015). However, our xCGE-LIF-based approach clearly distinguished Lc4 from its structural isomer Gb4 and even its linkage isomer nLc4 (Rossdam et al., 2019). In accordance to our observations, Lc4-Cer could be unambiguously detected from hESC- and hiPSC-derived GSLs by capillary liquid chromatography-based separation of structural isomers of tetraglycosylceramides and subsequent ESI/MS as well as by NMR analysis (Barone et al., 2013; Säljö et al., 2017), but these studies did not assess the effect of hPSC differentiation on Lc4-Cer levels.

We additionally performed flow cytometry using the BG-1 antibody specific for the T1LN structure (Domino et al., 2001) present in Lc4-Cer. To the best of our knowledge there is no antibody available which exclusively binds to Lc4-Cer. BG-1 will also detect *N*- and *O*-glycans containing the T1LN epitope which has been previously reported to be hPSC-specific by us and others (Hasehira et al., 2012; Konze et al., 2017; Natunen et al., 2011). Our flow cytometric analyses showed that the cell surface reactivity towards the BG-1 antibody strongly declined during differentiation, suggesting that not only Lc4-Cer but also levels of *N*- and *O*-glycans carrying T1LN structures decreased. Probably using the same antibody as we did (both are designated as clone K21), Liang *et al*. performed flow cytometry analyses and claimed a significant down-regulation of Lc4-Cer on hESCs upon differentiation into neural progenitor cells and definitive endoderm (Liang et al., 2011). However, according to the specificity of this antibody, they most likely also detected a decline of T1LN structures, irrespective to which type of glycan they belong to. Anyhow, while Liang *et al*. analyzed the cells after 2-3 weeks of differentiation, we differentiated the hESCs and hiPSCs for only 4 days into all three germ layers and thereby could show that T1LN already decreases in early stages of differentiation.

The β-1-3-specific transfer of galactose to either *N*-acetylglucosamine giving rise to T1LN structures (as present in the H-1 epitope detected by the SSEA-5 antibody or in Lc4-Cer) or to *N*-acetylgalactosamine as present in globoseries GSLs (e.g. SSEA-3, SSEA-4, globo H) is catalyzed by the enzyme β-1-3 galactosyltransferase (B3GALT5, (Zhou et al., 2000)). We noted in our study that differentiation is accompanied by rapid downregulation of *B3GALT5* expression. This finding is in accordance with previous studies showing a significant downregulation of *B3GALT5* gene expression upon hPSC differentiation (Liang et al., 2010; Liang et al., 2011; Ojima et al., 2015). Down-regulation of *B3GALT5* gene expression is likely to be associated with reduced synthesis of glycans containing β-1-3-linked galactose comprising glycans with T1LN structures. Interestingly, we observed that cell surface levels of glycans with terminal T1LN-structures decreased more rapidly upon differentiation than those of their fucosylated downstream products harboring a H-1 epitope (detected by the SSEA-5 antibody). Strikingly, the sorted T1LN-negative population on day 4 of differentiation still contained significant levels of glycans detected by the SSEA-5 antibody. In analogy to our findings, it was previously shown that hESCs or human embryonal carcinoma cells lose SSEA-3 expression before they lose expression of its biosynthetic product sialyl-SSEA-3 (SSEA-4) (Draper et al., 2002; Fenderson et al., 1987; Wright and Andrews, 2009). Based on these findings, we hypothesize that differentiation-induced down-regulation of *B3GALT5* will lead to a reduced synthesis of T1LN structures while existing T1LN structures are still further converted into their downstream products, e.g. H-1 or SSEA-4 antigens. Thus, during continuous cellular proliferation, levels of glycans with T1LN structures are more rapidly depleted from the cell surface than their biosynthetic products (Figure 4). This hypothesized mechanism can also explain the previously made observations for the SSEA3 and SSEA4 couple. Confirmatory to this model, we observed in the T1LN-negative population that SSEA3 levels were considerably lower than SSEA4 levels, but where interestingly still higher than T1LN levels. Thus, we claim that T1LN is a superior stem cell marker compared to SSEA-3, SSEA-4, and H-1 (fucosyl-Lc4 detected by the SSEA5 antibody) as it more rapidly declines upon onset of differentiation.

**Figure 4.**
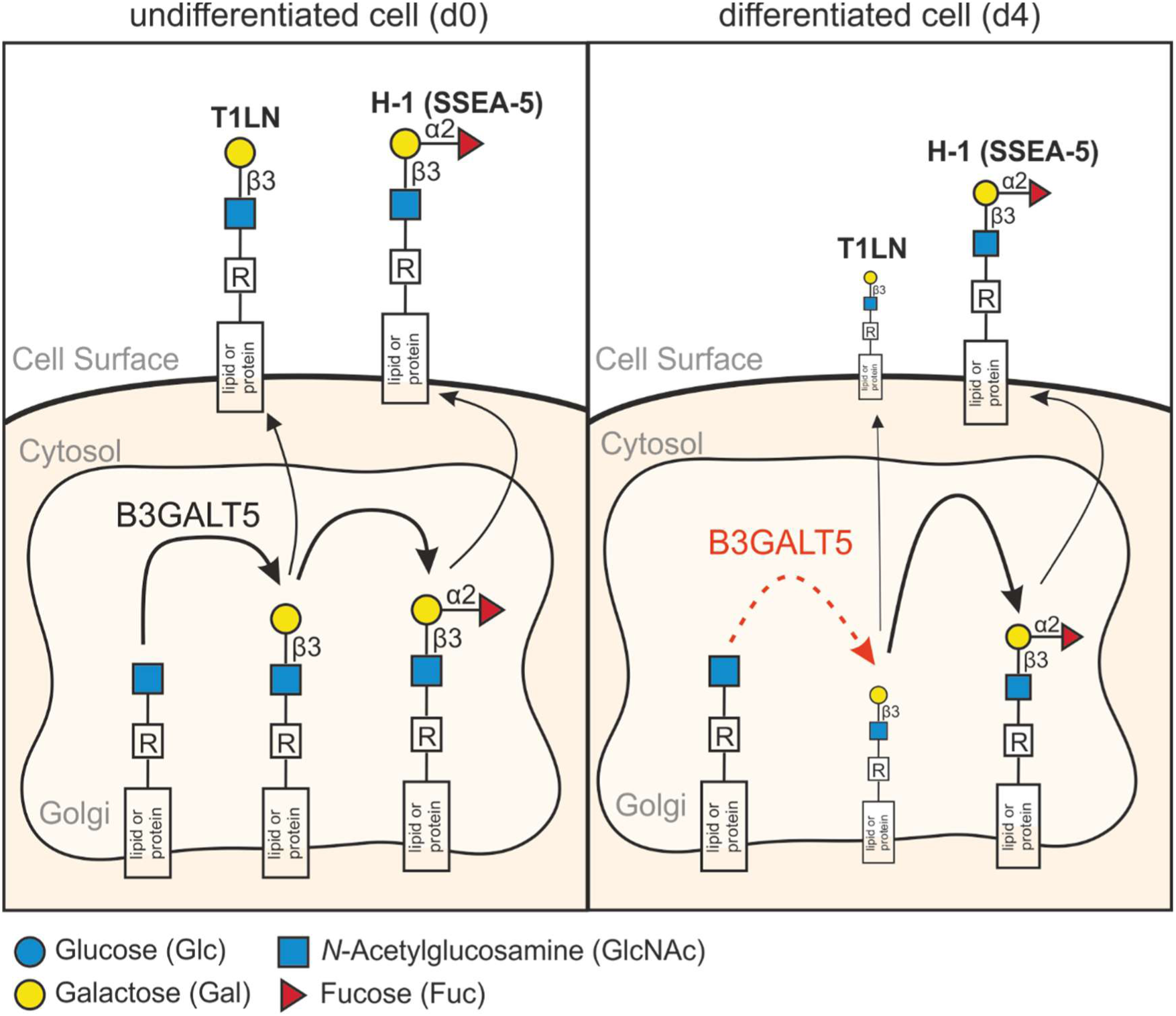
Model for T1LN expression during differentiation of hPSCs. In hPSCs (d0, left) B3GALT5 is highly expressed leading to continuous synthesis of protein and lipid-bound glycans decorated with T1LN structures that can be further elongated by fucosylation into H-1 epitopes which are target structures for the SSEA-5 antibody. During differentiation (e.g. on d4, right) expression of B3GALT4 drops and synthesis of glycans with T1LN epitopes is reduced while these glycans are still continuously fucosylated into H-1 structures. Therefore, levels of glycans with T1LN structures decrease more rapidly that H-1 glycans.

## Materials and Methods

### Human pluripotent stem cell culture

Cell culture reagents and cell culture vessels were obtained from Thermo Fisher Scientific (Waltham, MA) and Greiner (Greiner Bio-One, Frickenhausen, Germany), respectively, unless stated otherwise. Both the human embryonic stem cell (hESC) line (ES03: ES Cell International, National Stem Cell Bank Wisconsin, WI; H9: WA09 Wisconsin Alumni Research Foundation (Wicell Research Institute, Inc.) Madison, WI), as well as the human induced pluripotent stem cell (hiPSC) line (HSC_Iso4_ADCF-iPC2 “Phoenix” (Haase et al., 2017; Kempf et al., 2016a) were maintained at 37 °C with 5% CO_2_ and 85% relative humidity. Culture of hESCs was conducted under standard conditions on γ-irradiated human fibroblasts (human foreskin fibroblasts CCD919, ATCC, Manassas, VA) (Thiesler et al., 2016) in stem cell medium consisting of KnockOut™ DMEM supplemented with 20% KnockOut™ Serum Replacement, 1% (v/v) MEM Nonessential Amino Acids, 0.05% (v/v) GlutaMax™, 0.1 mM 2-mercaptoethanol (all reagents obtained from Life Technologies, Carlsbad, CA) and 50 ng/ml basic fibroblast growth factor (bFGF, Institute of Technical Chemistry, Leibniz University Hannover, Hannover, Germany). For passaging every 7 days, cells were incubated with 0.2% (w/v) Collagenase IV (Life Technologies) for 5-10 min at 37 °C and subsequently transferred onto new feeder cells, which were seeded 24 h prior. For further analysis, hESCs were cultivated at least three passages under feeder-free conditions on Matrigel^®^ (1:60 dilution, Corning, Corning, NY) in E8 medium (Chen et al., 2011) to exclude contaminating feeder cells. hiPSCs were routinely cultured under feeder-free conditions as monolayer on Matrigel^®^ in E8 medium (Rossdam et al., 2019). For passaging of both hiPSCs and hESCs every 3-4 days under feeder-free conditions, cells were incubated with 0.5 mM EDTA in PBS for 1 min at room temperature and subsequently reseeded onto new Matrigel^®^-coated flasks.

### Differentiation into the three germ layers

Differentiation into the three germ layers was performed in suspension culture in Corning^®^ Costar^®^ Ultra-Low Attachment 6-well-plates (Merck, Darmstadt, Germany). Prior to seeding, the cells were singularized using 0.5 mM EDTA in PBS for 10 min at 37 °C. For mesodermal/ ectodermal and endodermal differentiation, 3.3 x 10^5^ cells and 2.6 x 10^5^ cells per ml were seeded in 3 ml E8 medium supplemented with 10 µM Rho-associated coiled-coil kinase (ROCK) inhibitor Y-27632 (RI, STEMCELL Technologies, Vancouver, Canada), respectively. For both mesodermal and ectodermal differentiation, the medium was replaced 48 h after seeding on day −2 and again on day −1 of differentiation by E8 medium without RI supplementation. Mesodermal differentiation was performed as described previously by Kempf *et al*. with minor modifications (Kempf et al., 2016a). Briefly, the differentiation was initiated on d0 by replacing the medium with RPMI/B27™ minus insulin (RPMI 1640 supplemented with 1% (v/v) GlutaMax™, 2% (v/v) B27™ minus insulin) supplemented with 13.5 µM CHIR99021 (Selleckchem, Houston, TX). The medium was replaced on d1 with RPMI/B27™ minus insulin supplemented with 5 µM IWP-4 (STEMCELL Technologies). On d3 medium change was performed using RPMI/B27™ minus insulin without any supplementation. Ectodermal differentiation was adapted from a protocol previously described by Shi *et al*. (Shi et al., 2012). Briefly, the differentiation was initiated on d0 by replacing the E8 medium with neural induction medium (1:1 mixture of N-2 containing medium (DMEM/F12 medium (PAN Biotech, Aidenbach, Germany) supplemented with 1% (v/v) N-2 (Invitrogen), 5 µg/ml insulin (Merck), 1% (v/v) GlutaMax™, 1% (v/v) MEM Nonessential Amino Acids, 0.1 mM 2-mercaptoethanol and B27™ containing medium (Neurobasal medium (Invitrogen) supplemented with 2% (v/v) B27™ minus insulin and 1% (v/v) GlutaMax™) supplemented with SB431542 (10 µM) and Dorsomorphin (1 µM, both STEMCELL Technologies). Medium was replaced daily. Endodermal differentiation was performed as described previously by Diekmann *et al*. with minor modifications (Diekmann et al., 2015). Briefly, differentiation was initiated on d0 24 to 48 h after seeding by replacing the medium with RPMI/B27™ minus insulin (RPMI 1640 supplemented with 1 % (v/v) GlutaMax™, 1% (v/v) B27™ minus insulin) supplemented with 5 µM CHIR99021 and 33.3 ng Activin A (STEMCELL Technologies). Medium was exchanged daily without CHIR99021 supplementation. All differentiations were stopped on d4 of differentiation by harvesting the cells for further analyses.

### Glycolipid extraction and deglycosylation

Glycolipids were extracted from cells as previously described (Rossdam et al., 2019). Briefly, approximately 2 x 10^6^ cells were harvested on each day of differentiation and stored at −80 °C until further use. Glycolipids were extracted by homogenizing the cell pellets in 1:2 chloroform/methanol (v/v) by sonication. Cell debris and proteins were removed by centrifugation at 1620 x g for 5 min and the extract was transferred into a new glass vessel. To ensure a complete extraction, this step was repeated with 1:1 chloroform/methanol (v/v) and 2:1 chloroform/methanol (v/v), respectively. The pooled extracts were further desalted and purified on a Chromabond^®^ C_18_ ec poplyprolyene column (Macherey-Nagel, Düren, Germany). The extracted glycolipids were evaporated to dryness under nitrogen gas to remove organic solvents for deglycosylation. To release the glycan head groups from glycosphingolipids (GSLs), the extracted glycolipids were digested with the GSL-specific LudgerZyme ceramide glycanase (CGase) from *Hirudo medicinalis* (Ludger Ltd, Abingdon, UK) for 24 h at 37 °C in the provided reaction buffer (Ludger Ltd). Released glycans were fluorescently labeled with 8-aminopyrene-1,3,6-trisulfonic acid (APTS, Merck). Briefly, the glycans were mixed with 2 µl APTS (20 mM in 3.5 M citric acid), the reducing agent 2 µl 2-picoline borane complex (PB, 2 M in DMSO, Merck) and 2 µl water. Labeling was achieved by incubation for 16.5 h at 37 °C in the dark. The reaction was stopped by adding 100 µl 80:20 acetonitrile/water (v/v). The labeled glycans were purified by performing a HILIC-SPE as previously described (Rossdam et al., 2019). Briefly, the reaction mixture was applied onto a GHP membrane of a Pall Nanosep^®^ MF Centrifugal Device (pore size 0.45 µm, Merck) loaded with a 100 mg/ml Biogel P-10 Gel (Bio-Rad, Hercules, CA) solution. The excess of APTS and reducing agent were removed by washing the column with 80:20 acetonitrile/water (v/v) supplemented with and without 100 mM trimethylamine (pH 8.5), respectively. Labeled glycans were eluted in water and concentrated in a SpeedVac concentrator and stored at −20 °C until further use.

### Multiplexed capillary gel electrophoresis coupled to laser-induced fluorescence detection (xCGE-LIF)

Unless stated otherwise, xCGE-LIF equipment was purchased from Applied Biosystems™ (Thermo Fisher Scientific, Foster City, CA). xCGE-LIF of APTS-labelled glycosphingolipid-derived glycans was carried out as previously described (Rossdam et al., 2019). Briefly, APTS-labelled glycans were prepared for xCGE-LIF in a 1:10 dilution in water (water for chromatography, LC-MS grade, Merck) and mixed with 1 µl GeneScan™ 500 LIZ™ dye Size Standard (1:50 dilution in HiDi™ Formamide) in HiDi™ Formamide in a total volume of 10 µl in a MicroAmp™ Optical 96-Well Reaction Plate. The analysis was carried out for 30 min at 12 kV and 60 °C using an ABI PRISM^®^ 3100-*Avant* Genetic Analyzer (remodeled by advanced biolab service GmbH, Munich, Germany) equipped with a 4-Capillary Array of 50 cm length filled with POP-7™ Polymer. Data was further processed using the GeneMapper™ Software v3.7. The migration time range of 10 to 300 MTU of resulting xCGE-LIF electropherograms was further processed using Microsoft Word Excel and GraphPad Prism 5.01 software. Relative signal intensities were calculated for individual peak intensities (heights) in relation to the sum of all peak intensities and displayed as percentage enabling comparison of individual peaks from different samples. The in-house migration time data base (Rossdam et al., 2019) enabled annotation of obtained migration times to the target glycan structures Lc4, SSEA3, SSEA4, and SSEA5. Statistical analyses were performed using GraphPad Prism 5.01 software and *p*-values of <0.05 upon unpaired Student’s *t*-test were regarded as statistically significant and displayed.

### Flow cytometric analysis

Cell surface expression of T1LN structures, H-1 structures (detected by the anti-SSEA-5 antibody), and globoseries GSLs SSEA-3 and SSEA-4 was analyzed by flow cytometry on d0 and d4 of differentiation. For this, the cells were harvested and dissociated into single cells by incubation with Accutase (STEMCELL Technologies) for 10 min at 37 °C. To ensure complete dissociation, cells were filtered through a Pre-Separation Filter (50 µm, Sysmex, Norderstedt, Germany). For Pax-6 staining only, cells were fixed for 20 min at 4 °C with paraformaldehyde (PFA, 4 % (w/v) in PBS) and permeabilized with Tween-20 (0.1 % (v/v) in PBS) for 15 min at 4 °C. All cells were pelleted and resuspended in fresh FACS buffer (1 % (w/v) BSA and 1 mM EDTA in PBS) and approximately 3 x 10^5^ cells were incubated with the respective antibody or the isotype control for 20 min at 4 °C in dilutions as listed in Table S1. After antibody staining, the cells were washed and resuspended in FACS buffer and analyzed using a CyFlow ML flow cytometer (Sysmex Partec, Görlitz, Germany). The data were processed using FlowJo v7.6.4 software. Differentiation and stem cell marker expression was analyzed in the gated population of viable cells (determined in an unstained control of both d0 and d4). For quantification of cells positive for their respective differentiation marker, cells which showed intensities higher than 99% of the respective isotype control were regarded as positive for the respective marker. For quantification of cells positive for T1LN, SSEA-4, and H1-structures (SSEA-5) the mean fluorescence intensity (MFI) of the respective target was determined and subtracted from the MFI of the respective isotype control. Statistical analyses were performed using GraphPad Prism 5.01 software and *p*-values of <0.05 upon unpaired Student’s *t*-test were regarded as statistically significant.

### Magnetic activated cell sorting

To remove T1LN-positive cells from a mixed population, magnetic activated cell sorting (MACS) was applied using the mouse IgM BG-1 antibody (BioLegend, San Diego, CA) coupled to α-mouse IgM microbeads (Miltenyi Biotec, Bergisch Gladbach, Germany) (Table S1). Both hESCs and hiPSCs were differentiated into the three germ layers. On d0 and d4 of each differentiation, cells were harvested, dissociated, and filtered, as described above. Approximately 5 x 10^6^ cells of d0 and d4, respectively, were pooled in MACS buffer (0.5 % (w/v) BSA and 2 mM EDTA in PBS, degassed) and stained with mouse IgM BG-1 antibody for 20 min at 4 °C followed by staining with 20 µl α-mouse IgM microbeads in MACS buffer in a total volume of 100 µl for 15 min at 4 °C. Stained cells were washed twice with 1 ml PBS each and resuspended in 2 ml MACS buffer. MACS was performed applying an autoMACS^®^ Pro Separator using autoMACS^®^ columns (both Miltenyi Biotec) and the depletion program with a flow rate of 0.25 ml/min. Cell surface expression of T1LN, SSEA-3, SSEA-4, and H-1 (SSEA-5) was monitored pre-MACS in the d0-, d4-, and d0/d4-pooled population, as well as post-MACS in the T1LN-positive and T1LN-negative population using flow cytometry. The data were again processed using the FlowJo v7.6.4 software. Quantification of differentiation marker expression on d4 was determined as described above. For quantification of cells positive for T1LN, SSEA-3, SSEA-4, and H-1 (SSEA-5) viable cells were determined in an unstained control of d0 for all pre- and post-MACS cell populations after ensuring that sorting did not influence the set gate of viable cells. MFI values were again determined by the the FlowJo v7.6.4 software and subtracted from the MFI of the respective isotype control. Due to a limited number of cells, isotype controls were only carried out for the d0 and d4 population. Therefore, the MFI of the pooled and the post-MACS populations were subtracted from the mean MFI of the isotype controls of d0 and d4.

### Isolation of RNA

Approximately 10^6^ hESCs and hiPSCs were harvested on d0 and d4 of each differentiation, respectively. Additionally, 10^6^ cells of the d0/d4-pooled population and of the post-MACS T1LN-positive and T1LN-negative population were collected for RNA isolation, cDNA synthesis, and quantitative real-time PCR (qPCR) and stored at – 80 °C until further use. For RNA isolation, cells were resuspended in TRIzol (250 µl, Life Technologies). RNA was extracted by addition of 50 µl chloroform, short incubation, and centrifugation for 15 min at 11,000 x g. Isopropanol was added to the upper aqueous phase, followed by stirring, incubation for 10 min, and centrifugation for 10 min at 11,000 x g. The pellet was washed in 250 µl ethanol (70 % (v/v)) and centrifuged again for 5 min at 11,000 x g. Residual ethanol was removed and the extracted RNA pellet was resuspended in RNAse free water. Total RNA concentration was determined from absorption at 260 nm using an UV spectrophotometer (Implen, Munich, Germany). RNA was stored at – 80 °C until further use.

### cDNA synthesis

cDNA was synthesized from 2.5 µg extracted RNA per sample in a total reaction volume of 50 µl. Residual genomic DNA was removed by digestion with RQ1 DNase (2.5 U, Promega) for 30 min at 37 °C. The reaction was stopped by adding RQ1 DNase stop solution and incubating for 10 min at 70 °C. The cDNA was synthesized by using random hexamer primers (Thermo Fisher Scientific). Primers were annealed for 10 min at 25 °C followed by cDNA synthesis using Maxima H Minus Reverse Transcriptase (Thermo Fisher Scientific) and incubation for 30 min at 50 °C. The reaction was stopped by heating for 5 min to 85 °C. All reaction steps were performed in a Piko PCR cycler (Thermo Fisher Scientific). Synthesized cDNA was diluted in water to 5 ng/µl and stored at – 20 °C.

### Quantitative real-time PCR (RT-qPCR)

RT-qPCR was performed in a total reaction volume of 10 µl containing 15 ng cDNA, 2 pmol forward/ reverse primer pairs (sequences listed in Table S2), and the BIO SyGreen Lo-ROX mix (PCR Biosystems, London, UK) on an ImageQuantQ3 Real-time PCR System (Thermo Fisher Scientific) in sealed 96-well optical reaction plates (Applied Biosystems). The specificity of the designed primers was ensured by melting-curve analysis and sequencing of the amplicon. The PCR was run for 40 two-step cycles (5 s at 95 °C, 30 s at 60 °C) after initial denaturation for 2 min at 50 °C and 3 min at 95 °C. Amplicon purity was ensured by melting-curve analysis in an additional cycle (15 s at 95 °C, 60 s at 60 °C, 15 s at 95 °C). Relative expression of target genes was determined by normalization to house-keeping genes (listed in Table S2). The ImageQuant software calculated cycle threshold (Ct) values automatically and the relative expression of target genes were subsequently calculated by determination of 2^-ΔΔCt^-values. Statistical analyses were performed using GraphPad Prism 5.01 software and *p*-values of <0.05 upon unpaired Student’s *t*-test were regarded as statistically significant.

## Supplementary data

Supplementary data for this article are available at *Glycobiology* online.

## Author contributions

C.R., S.B., J.B., A.O., M.D.A, and O.N. conducted the experiments, C.R. and F.F.R.B. designed the experiments and performed data analysis and interpretation. F.F.R.B. conceived and supervised this study. C.R. and F.F.R.B. wrote the manuscript, and all authors reviewed, edited and agreed on its contents.

## Acknowledgments

The authors would like to thank Prof. Dr. Gerardy-Schahn and Prof. Dr. Christoph Garbers, Institute of Clinical Biochemistry, Hannover Medical School (MHH) for providing general laboratory equipment.

## Supplementary information

**Table S1.** Antibodies used for flow cytometry and magnetic activated cell sorting.

| Primary antibodies and conjugated antibodies |  |  |  |  |  |
| --- | --- | --- | --- | --- | --- |
| Target | Host species (isotype) | Supplier | Order # | Dilution | Conjugated fluorochrome |
| CXCR4 (CD184) | recombinant human IgG1 | Miltenyi Biotec | 130-117-504 | 1:50 | PE |
| EpCAM (CD326) | recombinant human IgG1 | Miltenyi Biotec | 130-111-116 | 1:50 | PE |
| IgG1 control | mouse IgG1, $\kappa$ | Miltenyi Biotec | 130-113-196 | 1:50 | APC |
| IgG1 control | recombinant human IgG1 | Miltenyi Biotec | 130-120-709 | 1:50 | APC |
| IgG1 control | recombinant human IgG1 | Miltenyi Biotec | 130-104-613 | 1:50 | PE |
| IgG<sub>2b</sub> control | mouse IgG<sub>2b</sub> | BD Biosciences | 565381 | 1:100 | APC |
| IgM control | mouse IgM, $\kappa$ | Miltenyi Biotec | 130-120-070 | 1:50 | PE |
| BG1 | mouse IgM, $\kappa$ | Biolegend | 932102 | 1:100 | / |
| NCAM (CD56) | mouse IgG<sub>2b</sub> | BD Biosciences | 341027 | 1:100 | APC |
| Pax-6<sup>1</sup> | rabbit IgG | BioLegend | 901301 | 1:50 | / |
| Pax-6 | rabbit IgG | Proteintech | 12323-1-AP | 1:100 | / |
| SSEA-3 | Rat IgM | Thermo Fisher Scientific | MA1-020-D650 | 1:100 | DyLight 650 |
| SSEA-4 | recombinant human IgG1 | Miltenyi Biotec | 130-098-347 | 1:10 | APC |
| SSEA-5 | mouse IgG1, $\kappa$ | Miltenyi Biotec | 130-106-661 | 1:10 | APC |
| Secondary antibodies and conjugated microbeads |  |  |  |  |  |
| Target | Host species (isotype) | Supplier | Order # | Dilution | Fluorochrome / microbeads |
| mouse IgM | rat | BD Biosciences | 553409 | 1:50 | PE |
| mouse IgM | rat | Miltenyi Biotec | 130-047-302 | 1:5 | microbeads |
| rabbit IgG | goat | Molecular Probes | A21245 | 1:1200 | Alexa 647 |

<sup>†</sup>The anti-Pax-6 antibody (901301) from BioLegend was unavailable for a long period due to difficulties in shipping. Therefore, for experiments shown in Fig. 2, Fig 3, as well as SFig. 2, and SFig 4 the anti-Pax-6 antibody (12323-1-AP) from Proteintech was used.

**Table S2.**
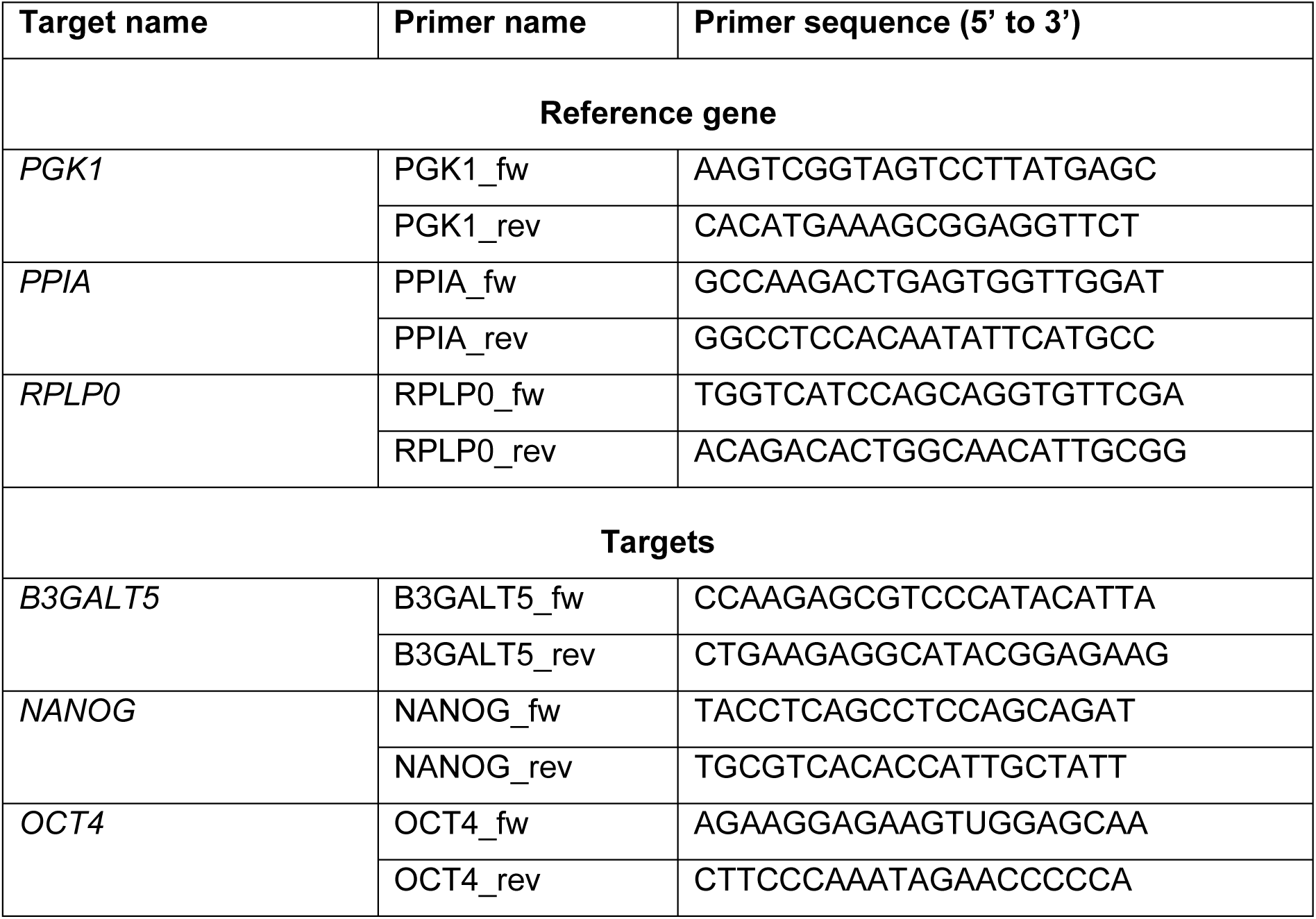
Oligonucleotides used as primers for qPCR.

**Supplemental Figure 1.**
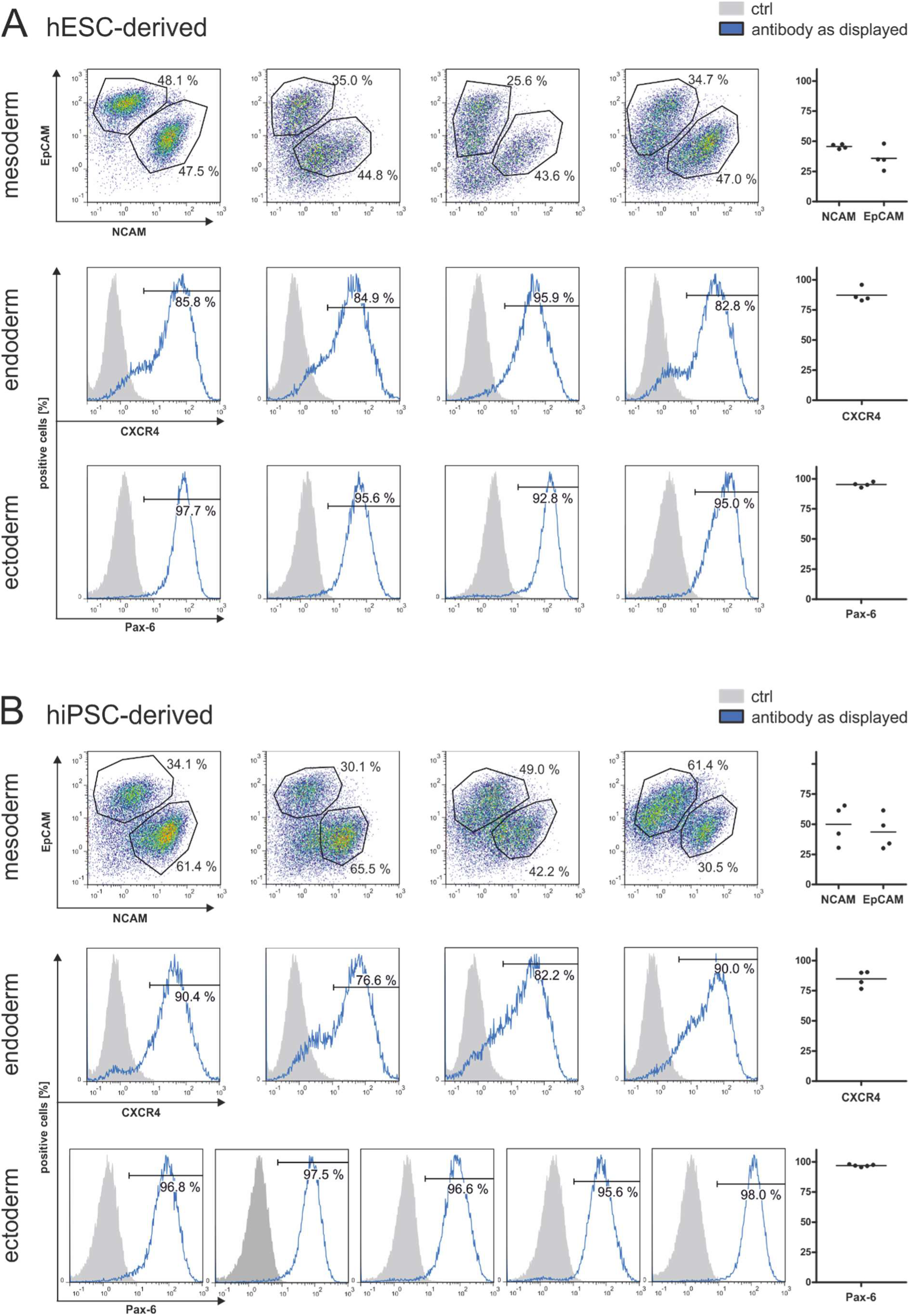
Quality control of differentiation approaches of replicates shown in Figure 1. (A-B) Differentiation of hESCs (A) and hiPSCs (B) into mesoderm (top panel), endoderm (middle panel), and ectoderm (bottom panel) was confirmed by cell surface staining of corresponding differentiation markers and quantified via flow cytometry. For mesoderm, dot plots are displayed showing balanced NCAM and EpCAM expression in viable cells on d4 of differentiation. For endoderm and ectoderm, overlay histograms show fluorescence signal of viable cells on d4 of differentiation (blue) for CXCR4 and Pax-6, respectively. Cells were defined positive if the fluorescence signal was above that of 99 % of the cells stained with the respective control (ctrl, grey). Scatter plots display all analyzed replicates shown in Figure 1.

**Supplemental Figure 2.**
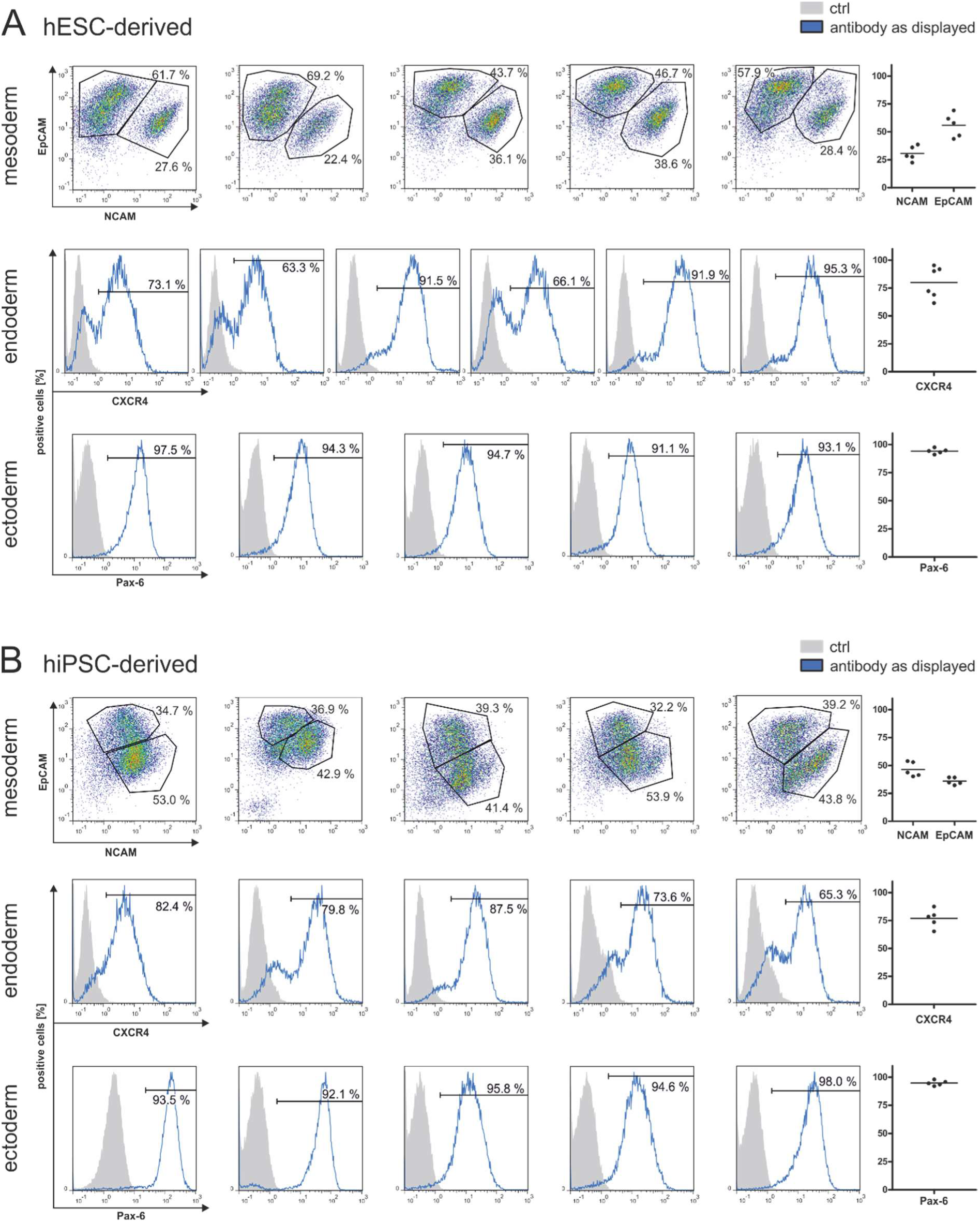
Quality control of differentiation approaches of replicates shown in Figure 2. (A-B) Differentiation of hESCs (A) and hiPSCs (B) into mesoderm (top panel), endoderm (middle panel), and ectoderm (bottom panel) was confirmed by cell surface staining of corresponding differentiation markers and quantified via flow cytometry. For mesoderm, dot plots are displayed showing balanced NCAM and EpCAM expression in viable cells on d4 of differentiation. For endoderm and ectoderm, overlay histograms show fluorescence signal of viable cells on d4 of differentiation (blue) for CXCR4 and Pax-6, respectively. Cells were defined positive if the fluorescence signal was above that of 99 % of the cells stained with the respective control (ctrl, grey). Scatter plots display all analyzed replicates shown in Figure 2.

**Supplemental Figure 3.**
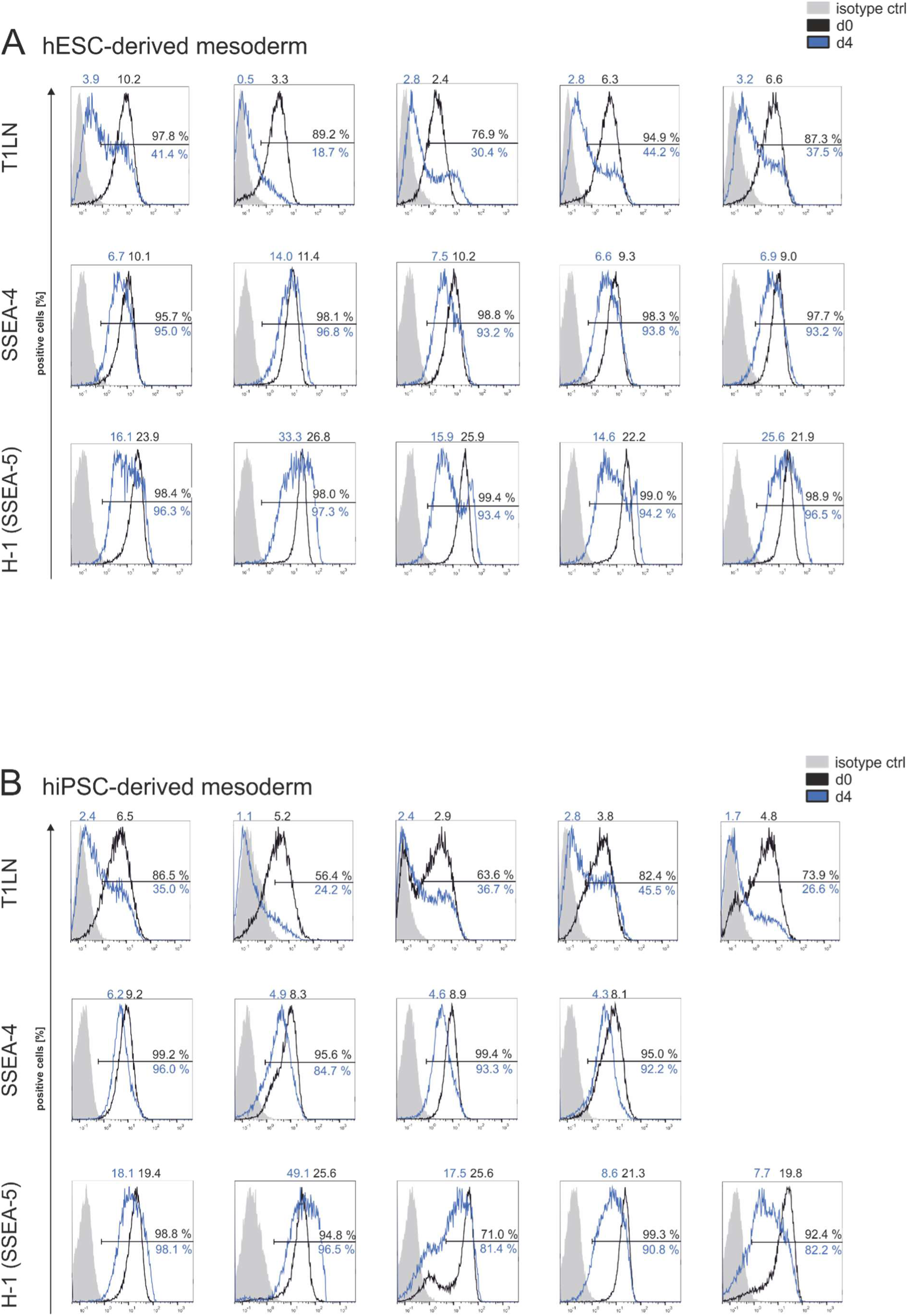
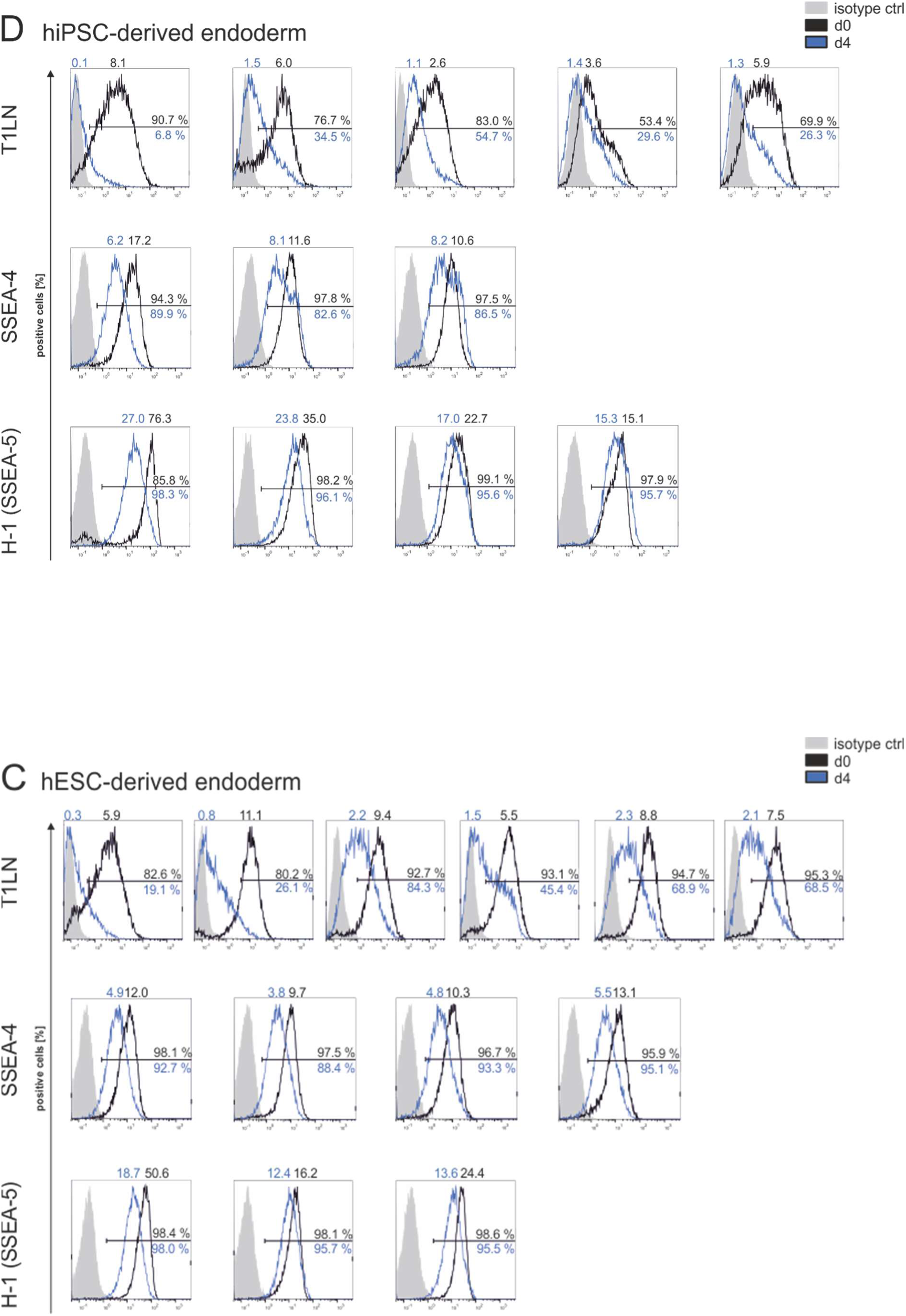
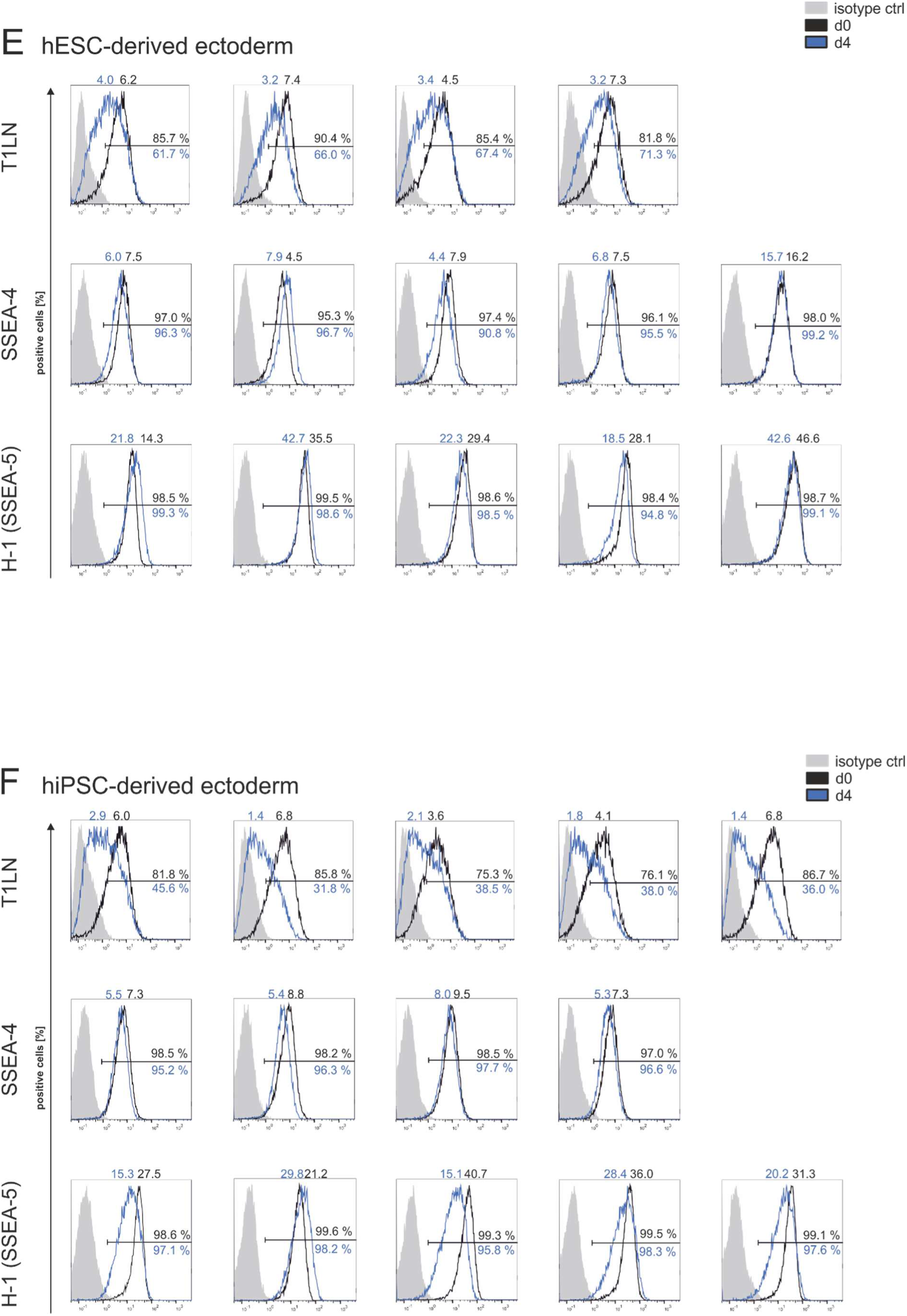
Cell surface expression of T1LN, SSEA-4, and SSEA-5 in hPSCs and differentiated derivatives of all replicates shown in Figure 2. (A-F) Flow cytometry analysis of hESC-(A) and hiPSC-derived mesoderm (B), hESC-(C) and hiPSC-derived endoderm (D), and hESC-(E) and hiPSC-derived ectoderm (F) on d0 and d4 of differentiation for T1LN (top), SSEA-4 (middle), and SSEA-5 (bottom). Overlay histograms show fluorescence signal of viable cells on d0 (black) and d4 (blue). Cells were defined positive if the fluorescence signal was above that of 99 % of the cells stained with the respective isotype control (grey). Only isotype controls for d0 are displayed but for calculation of positive cells on d4, isotype controls of d4 were considered. Number of positive cells are shown below (blue) and above (black) the line representing the gate. Mean fluorescence intensity is shown above histograms (d0, black and d4, blue).

**Supplemental Figure 4.**
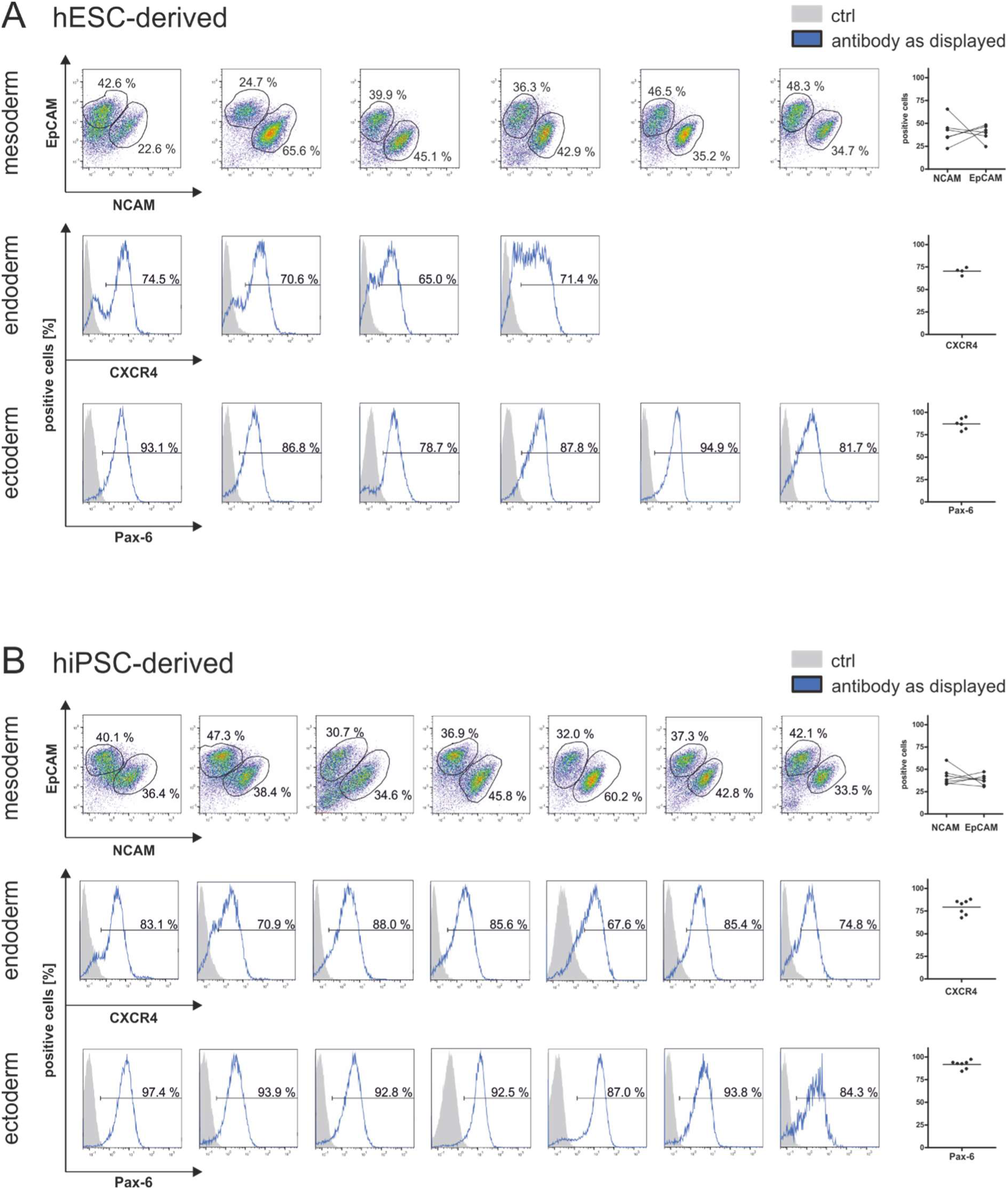
Quality control of differentiation approaches of replicates shown in Figure 3. (A-B) Differentiation of hESCs (A) and hiPSCs (B) into mesoderm (top panel), endoderm (middle panel), and ectoderm (bottom panel) was confirmed by cell surface staining of corresponding differentiation markers and quantified via flow cytometry. For mesoderm, dot plots are displayed showing balanced NCAM and EpCAM expression in viable cells on d4 of differentiation. For endoderm and ectoderm, overlay histograms show fluorescence signal of viable cells on d4 of differentiation (blue) for CXCR4 and Pax-6, respectively. Cells were defined positive if the fluorescence signal was above that of 99 % of the cells stained with the respective control (ctrl, grey). Scatter plots display all analyzed replicates shown in Figure 3.

**Supplemental Figure 5.**
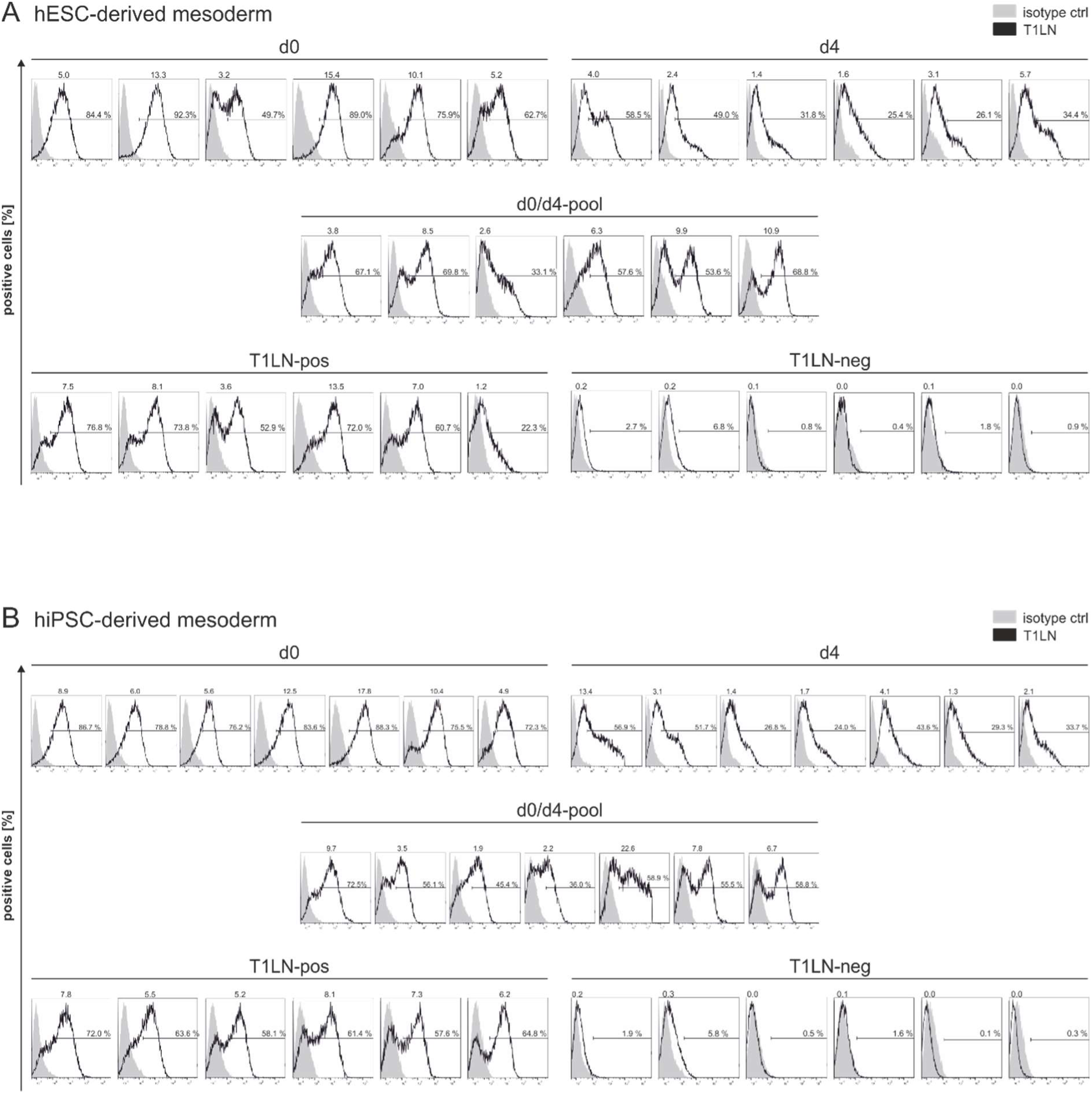
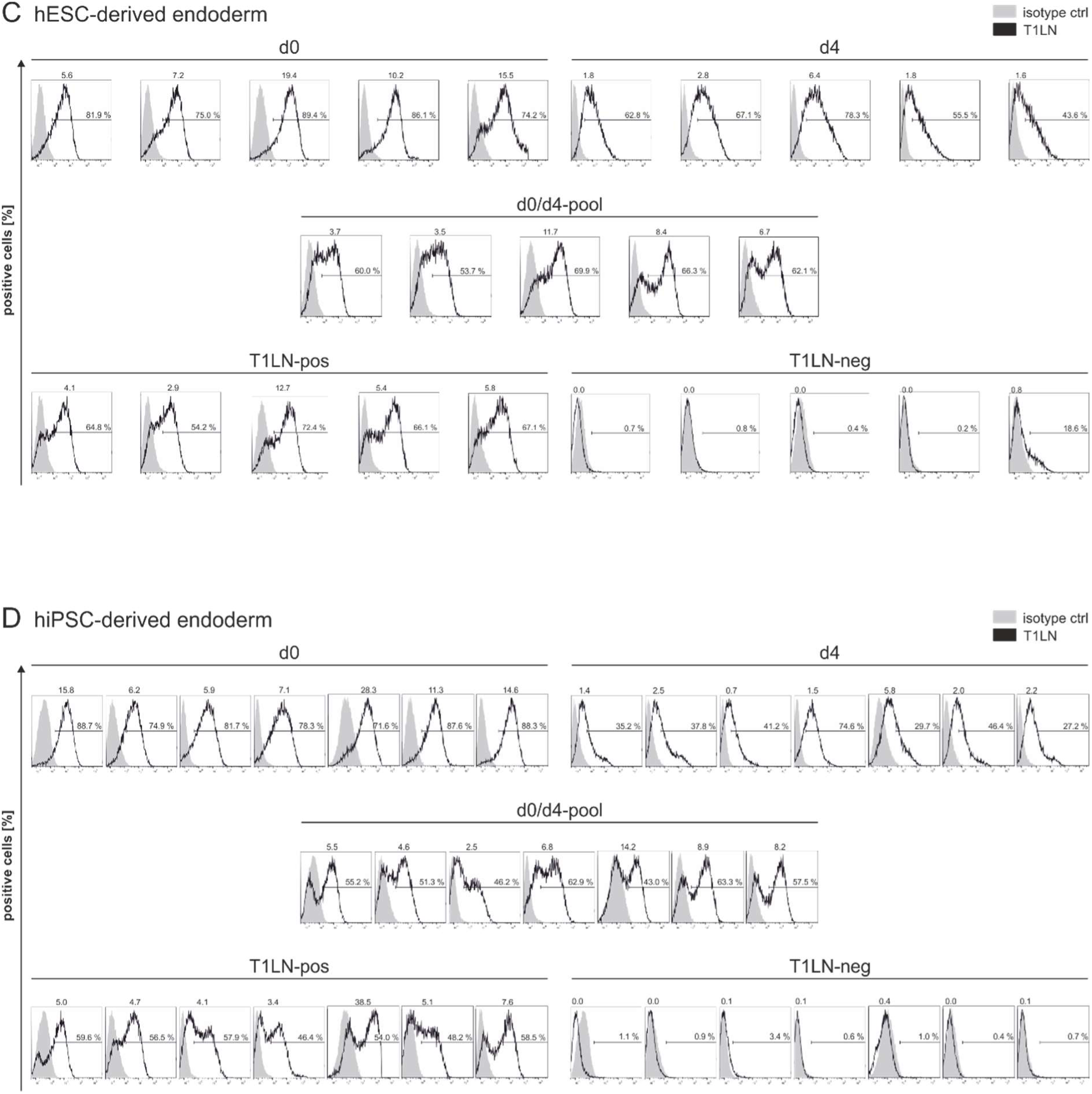
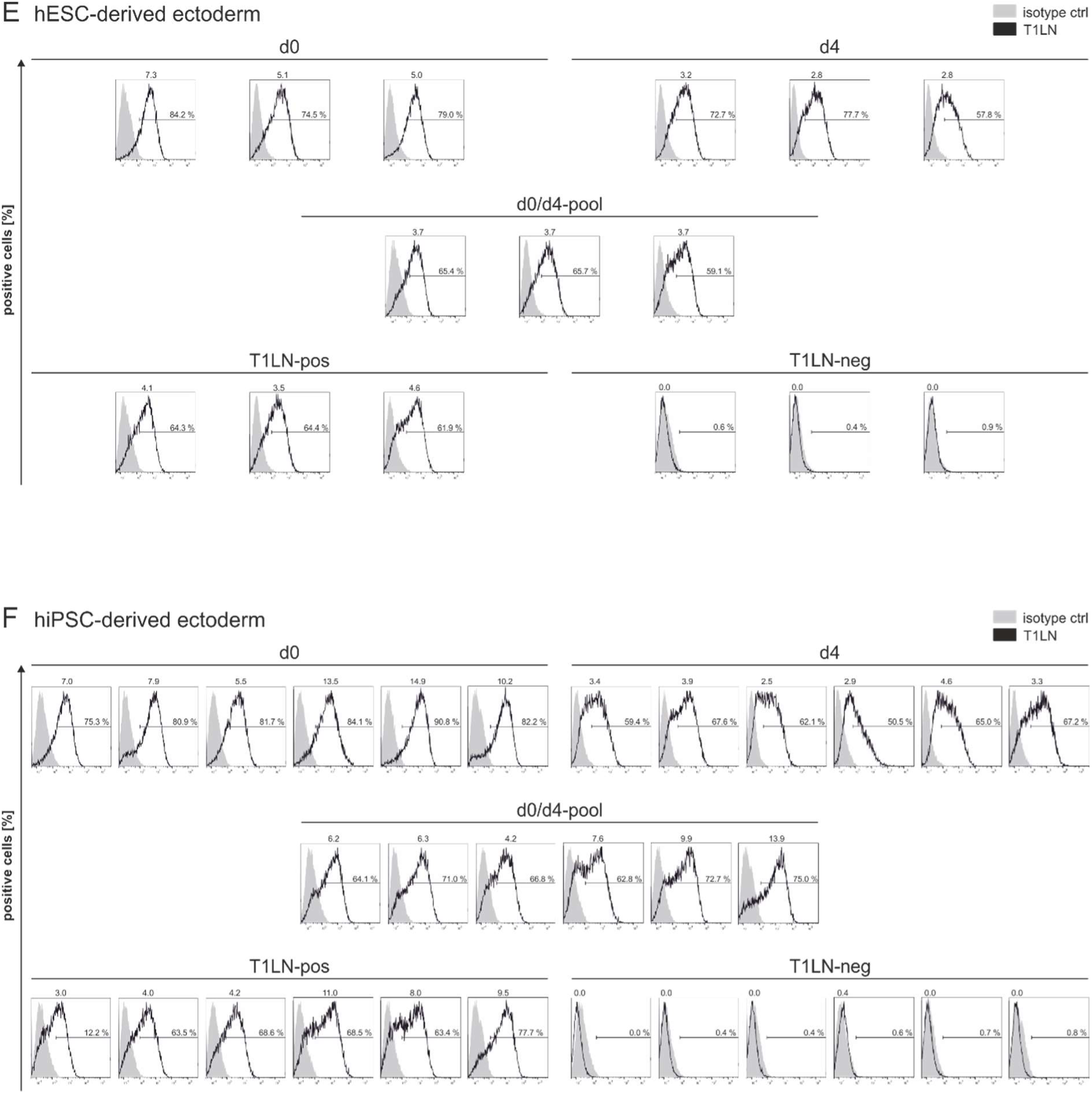
Cell surface expression of T1LN in hPSCs and differentiated derivatives pre- and post-MACS of all replicates shown in Figure 3. (A-E) Cell surface expression was analyzed using flow cytometry pre-MACS in d0 (top left), d4 (top right) and d0/d4-pooled cells (middle), as well as post-MACS in the T1LN-positive (bottom left) and T1LN-negative population (bottom right) in hESC-(A) and hiPSC-derived mesoderm (B), hESC-(C) and hiPSC-derived endoderm (D), and hESC-(E) and hiPSC-derived ectoderm (F). Overlay histograms show fluorescence signal of viable cells (black) in regard to the respective isotype control (isotype ctrl, shown in grey). Cells were defined positive if the fluorescence signal was above that of 99 % of the cells stained with the respective isotype control. Only isotype controls for d0 are displayed, but for calculation of positive cells on d4 isotype controls of d4 were considered. For calculation of positive cells post-MACS, both isotype controls of d0 and d4 were considered. Number of positive cells are shown above the line representing the gate. Mean fluorescence intensity is shown above histograms.

**Supplemental Figure 6.**
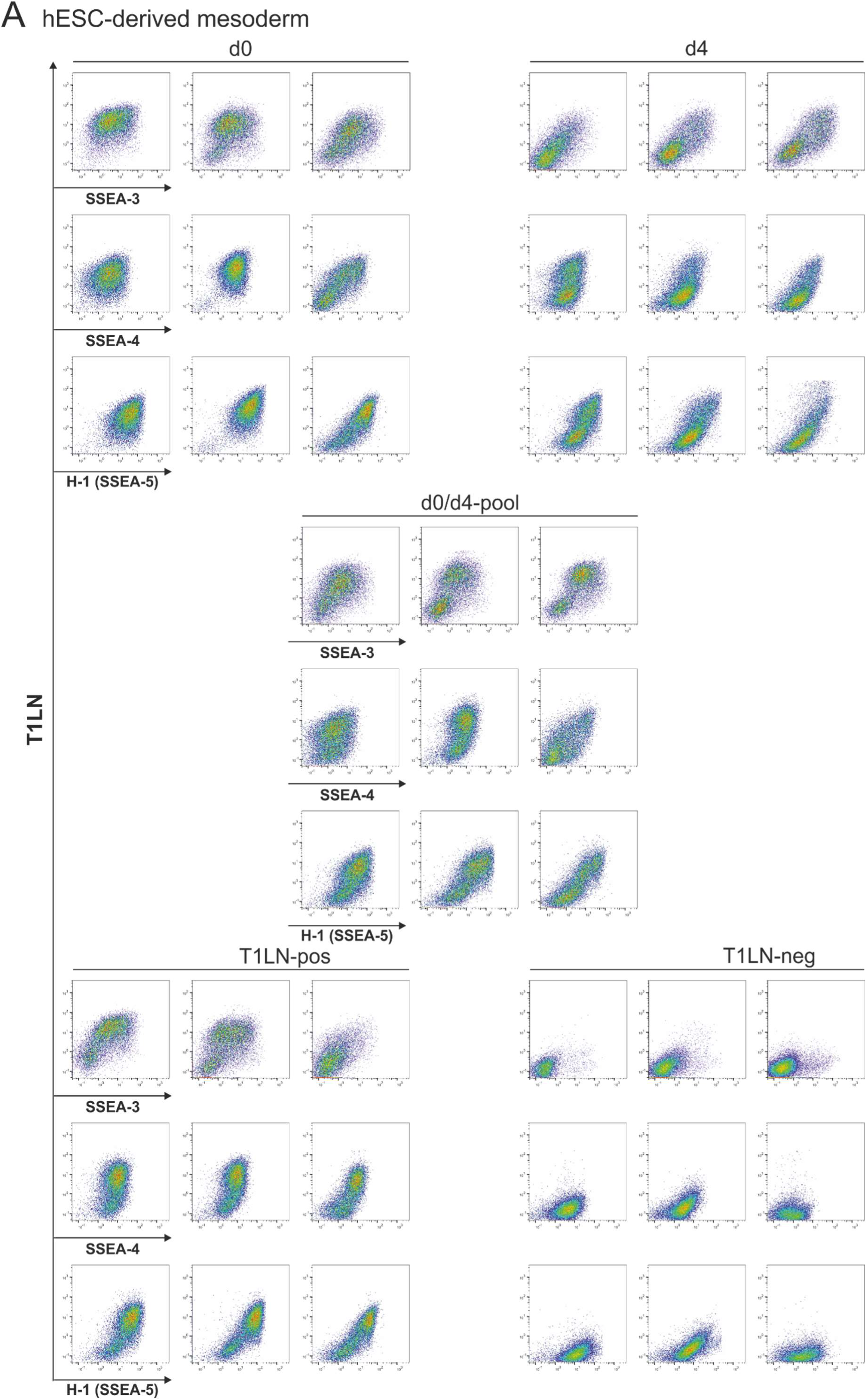
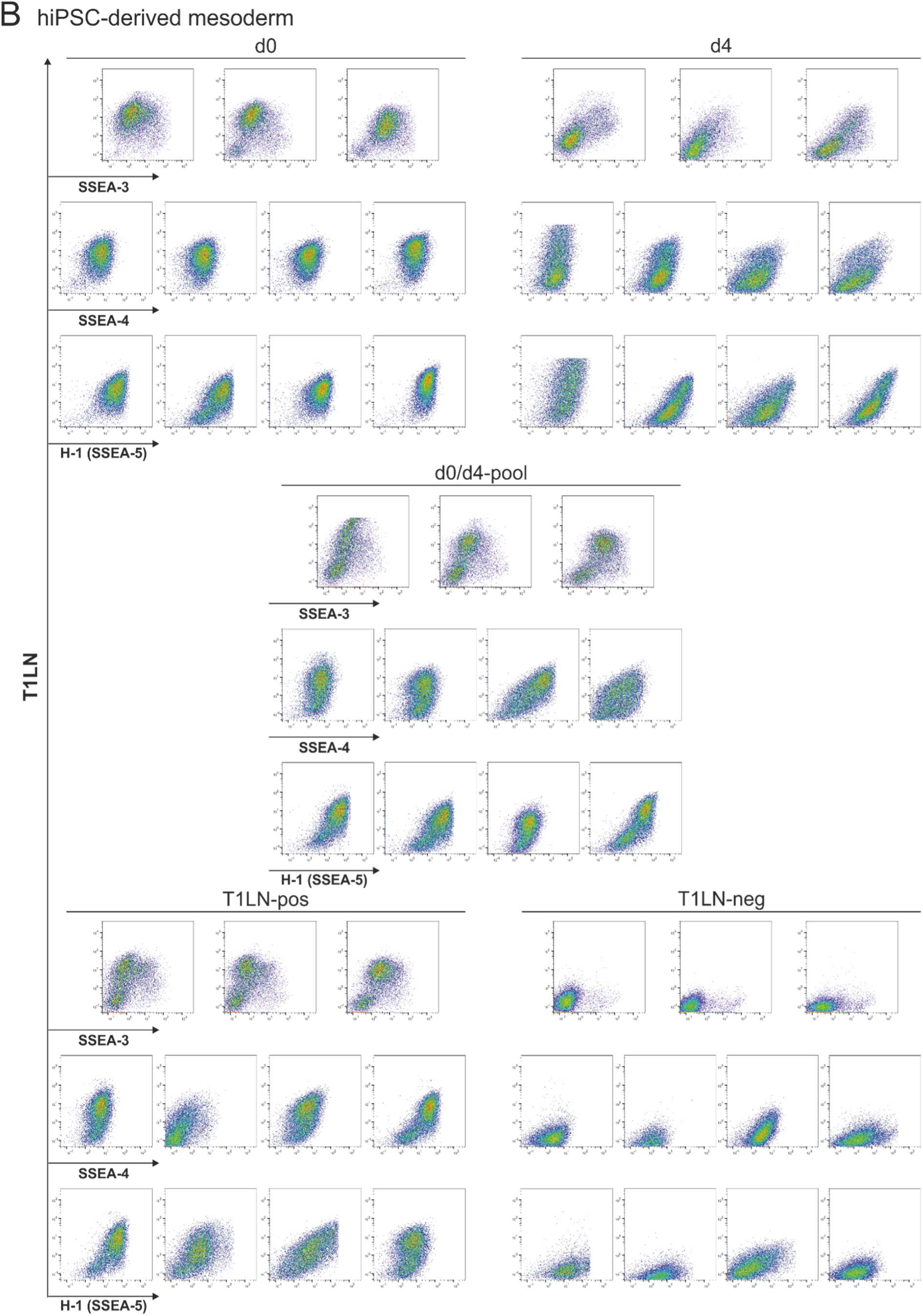
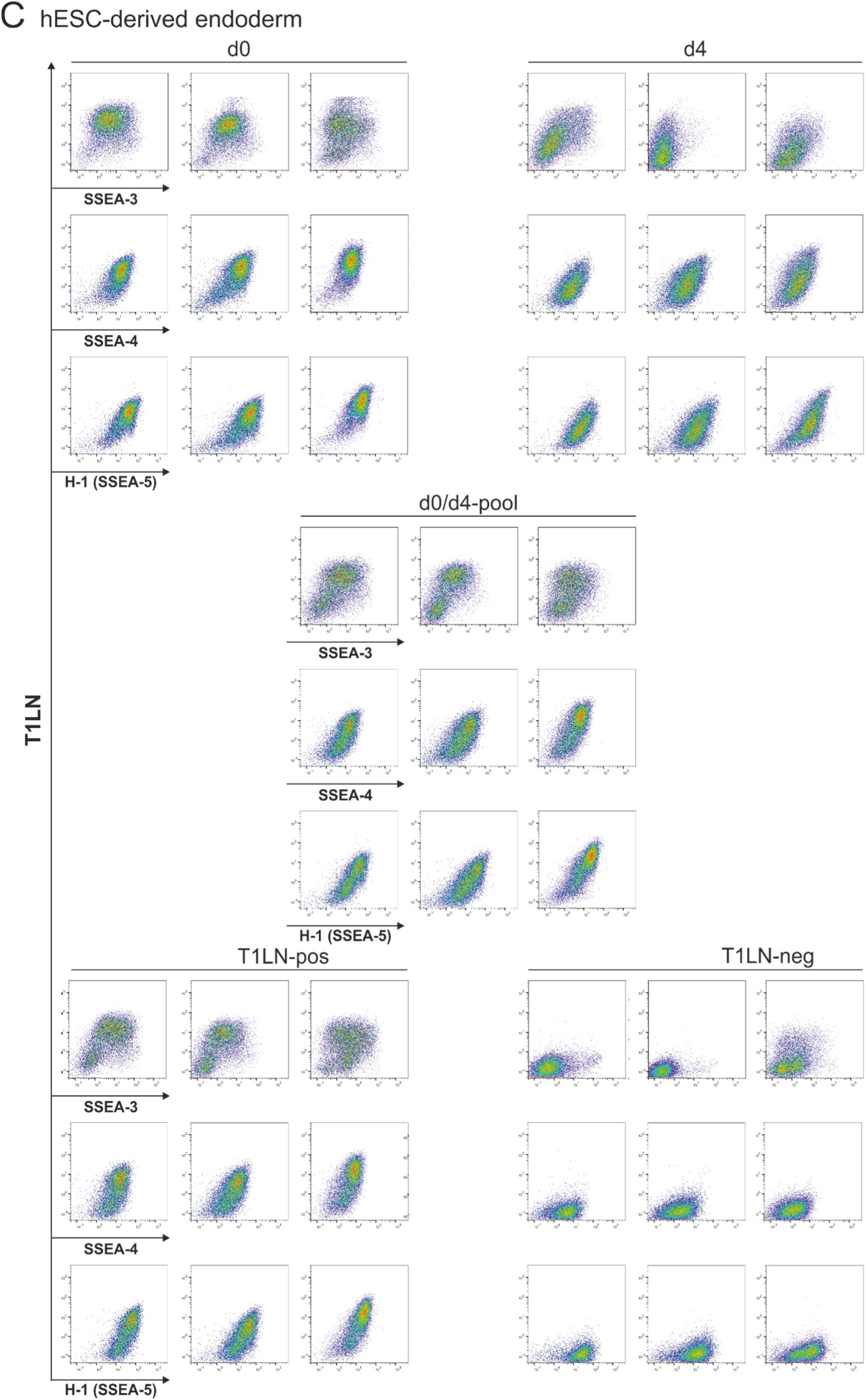
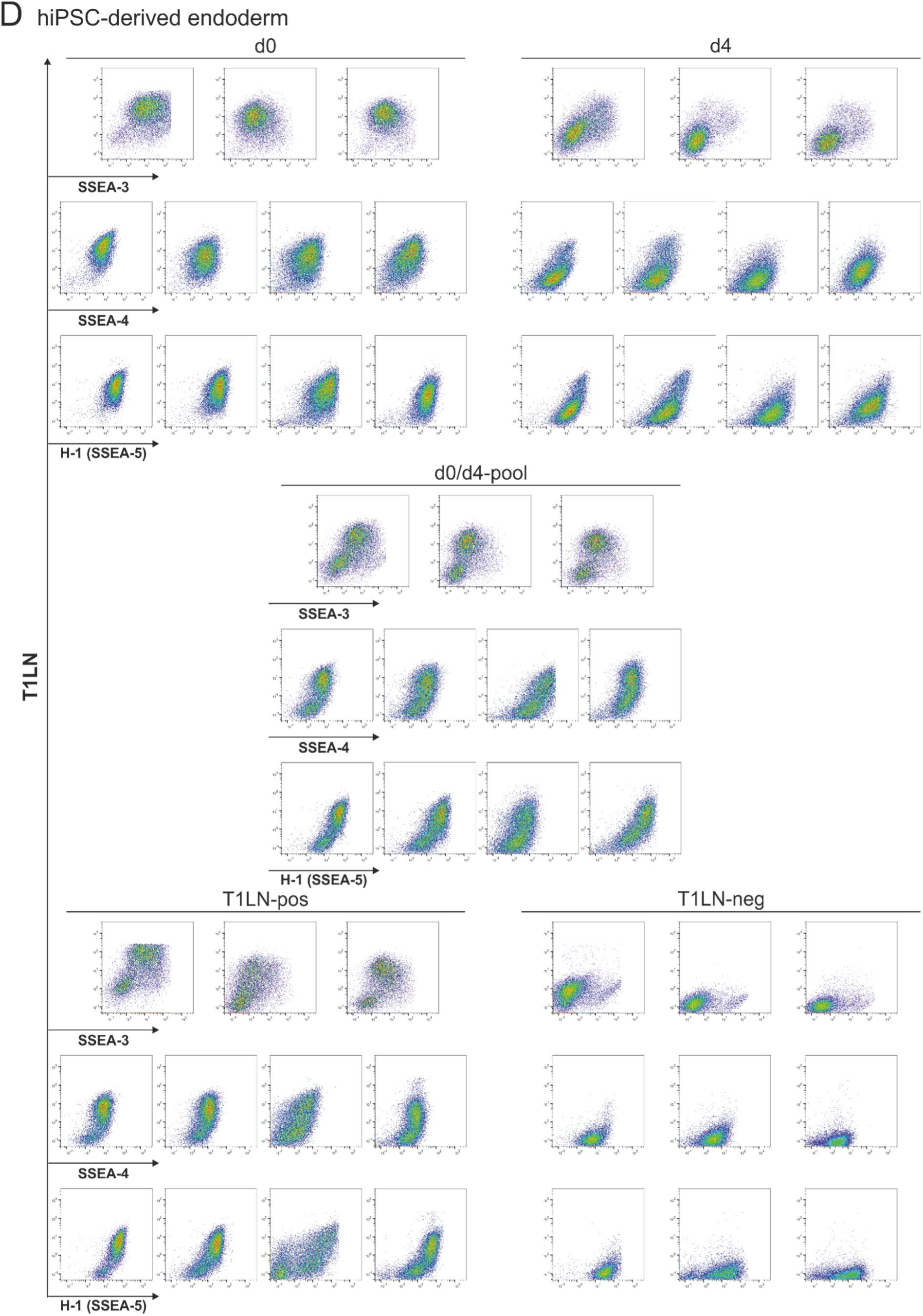
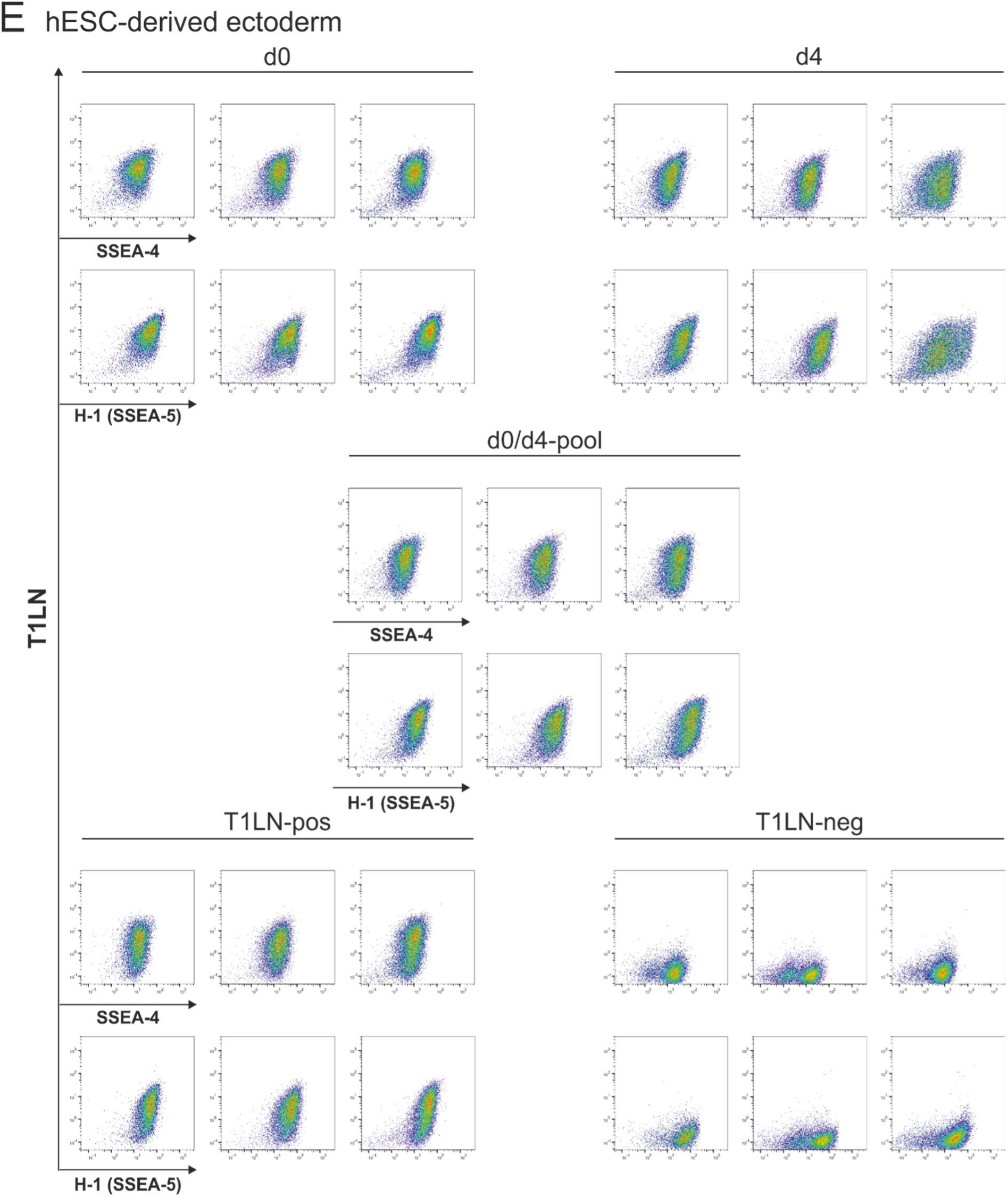
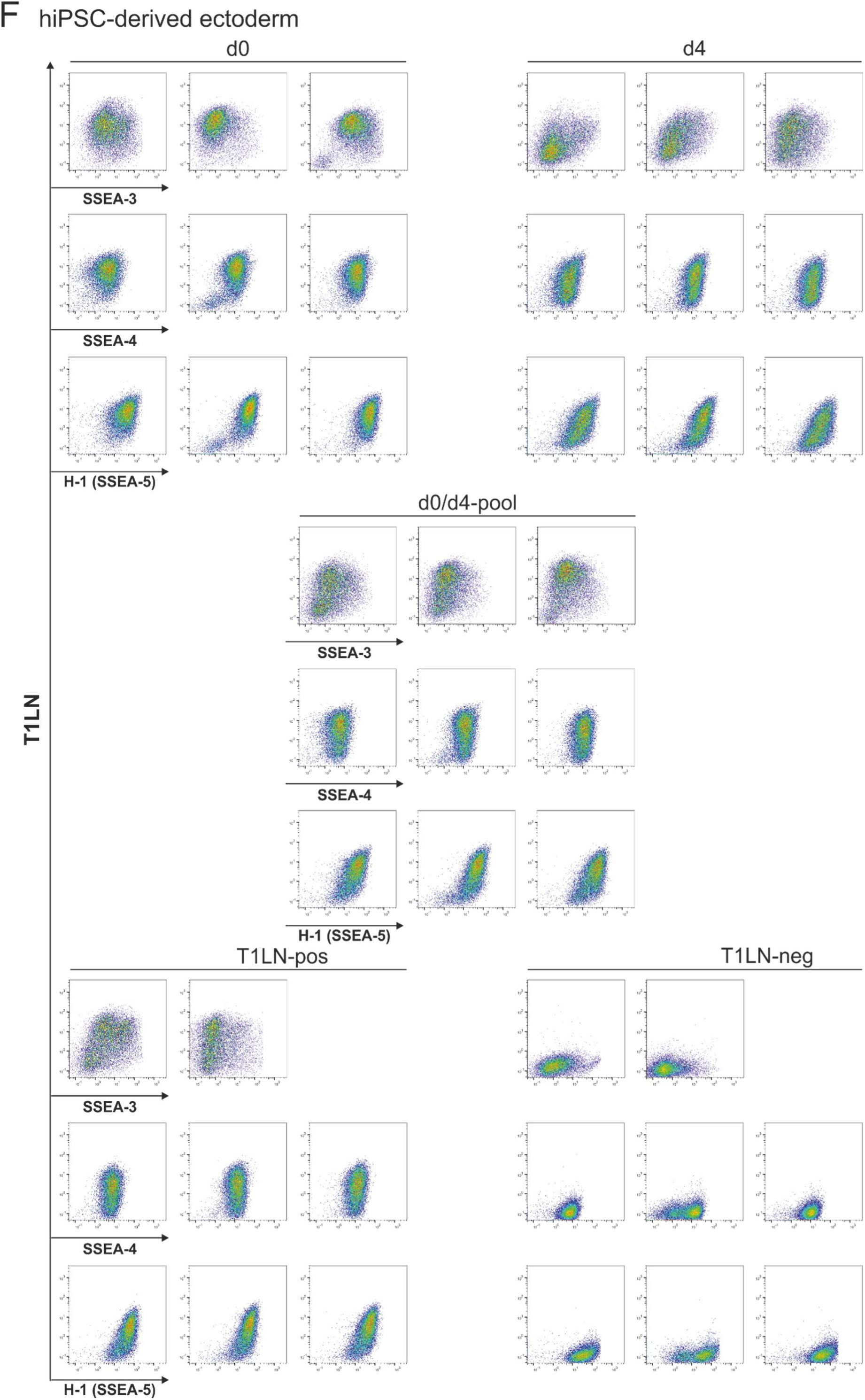
Cell surface expression of T1LN, SSEA-3, SSEA-4, and SSEA-5 in hPSCs and differentiated derivatives pre- and post-MACS of all replicates shown in Figure 3. (A-F) Cell surface expression was analyzed using flow cytometry pre-MACS in d0 (top left), d4 (top right) and d0/d4-pooled cells (middle), as well as post-MACS in the T1LN-positive (bottom left) and T1LN-negative population (bottom right) in hESC-(A) and hiPSC-derived mesoderm (B), hESC-(C) and hiPSC-derived endoderm (D), and hESC-(E) and hiPSC-derived ectoderm (F). Dot plots display cell surface expression of T1LN and SSEA-3 (top panel), Lc4 and SSEA-4 (middle panel), as well as cell surface expression of T1LN and SSEA-5 (bottom panel).

## References

Barone, A., Benktander, J., Ångström, J., Aspegren, A., Björquist, P., Teneberg, S., and Breimer, M.E. (2013). Structural complexity of non-acid glycosphingolipids in human embryonic stem cells grown under feeder-free conditions. J Biol Chem 288, 10035–10050. 10.1074/jbc.M112.436162.

Barone, A., Säljö, K., Benktander, J., Blomqvist, M., Månsson, J.E., Johansson, B.R., Mölne, J., Aspegren, A., Björquist, P., Breimer, M.E., and Teneberg, S. (2014). Sialyl-lactotetra, a novel cell surface marker of undifferentiated human pluripotent stem cells. J Biol Chem 289, 18846–18859. 10.1074/jbc.M114.568832.

Carpenter, M.K., Rosler, E.S., Fisk, G.J., Brandenberger, R., Ares, X., Miura, T., Lucero, M., and Rao, M.S. (2004). Properties of four human embryonic stem cell lines maintained in a feeder-free culture system. Dev Dyn 229, 243–258. 10.1002/dvdy.10431.

Chen, G., Gulbranson, D.R., Hou, Z., Bolin, J.M., Ruotti, V., Probasco, M.D., Smuga-Otto, K., Howden, S.E., Diol, N.R., Propson, N.E., et al. (2011). Chemically defined conditions for human iPSC derivation and culture. Nat Methods 8, 424–429. 10.1038/nmeth.1593.

Choo, A.B., Tan, H.L., Ang, S.N., Fong, W.J., Chin, A., Lo, J., Zheng, L., Hentze, H., Philp, R.J., Oh, S.K., and Yap, M. (2008). Selection against undifferentiated human embryonic stem cells by a cytotoxic antibody recognizing podocalyxin-like protein-1. Stem Cells 26, 1454–1463. 10.1634/stemcells.2007-0576.

Cumin, C., Huang, Y.L., Rossdam, C., Ruoff, F., Céspedes, S.P., Liang, C.Y., Lombardo, F.C., Coelho, R., Rimmer, N., Konantz, M., et al. (2022). Glycosphingolipids are mediators of cancer plasticity through independent signaling pathways. Cell Rep 40, 111181. 10.1016/j.celrep.2022.111181.

D’Angelo, G., Capasso, S., Sticco, L., and Russo, D. (2013). Glycosphingolipids: synthesis and functions. Febs j 280, 6338–6353. 10.1111/febs.12559.

Diekmann, U., Lenzen, S., and Naujok, O. (2015). A reliable and efficient protocol for human pluripotent stem cell differentiation into the definitive endoderm based on dispersed single cells. Stem Cells Dev 24, 190–204. 10.1089/scd.2014.0143.

Domino, S.E., Zhang, L., Gillespie, P.J., Saunders, T.L., and Lowe, J.B. (2001). Deficiency of reproductive tract alpha(1,2)fucosylated glycans and normal fertility in mice with targeted deletions of the FUT1 or FUT2 alpha(1,2)fucosyltransferase locus. Mol Cell Biol 21, 8336–8345. 10.1128/mcb.21.24.8336-8345.2001.

Draper, J.S., Pigott, C., Thomson, J.A., and Andrews, P.W. (2002). Surface antigens of human embryonic stem cells: changes upon differentiation in culture. J Anat 200, 249–258. 10.1046/j.1469-7580.2002.00030.x.

Fenderson, B.A., Andrews, P.W., Nudelman, E., Clausen, H., and Hakomori, S. (1987). Glycolipid core structure switching from globo- to lacto- and ganglio-series during retinoic acid-induced differentiation of TERA-2-derived human embryonal carcinoma cells. Dev Biol 122, 21–34. 10.1016/0012-1606(87)90328-9.

Fong, C.Y., Gauthaman, K., and Bongso, A. (2010). Teratomas from pluripotent stem cells: A clinical hurdle. J Cell Biochem 111, 769–781. 10.1002/jcb.22775.

Fujitani, N., Furukawa, J., Araki, K., Fujioka, T., Takegawa, Y., Piao, J., Nishioka, T., Tamura, T., Nikaido, T., Ito, M., et al. (2013). Total cellular glycomics allows characterizing cells and streamlining the discovery process for cellular biomarkers. Proc Natl Acad Sci U S A 110, 2105–2110. 10.1073/pnas.1214233110.

Haase, A., Göhring, G., and Martin, U. (2017). Generation of non-transgenic iPS cells from human cord blood CD34(+) cells under animal component-free conditions. Stem Cell Res 21, 71–73. 10.1016/j.scr.2017.03.022.

Hasehira, K., Tateno, H., Onuma, Y., Ito, Y., Asashima, M., and Hirabayashi, J. (2012). Structural and quantitative evidence for dynamic glycome shift on production of induced pluripotent stem cells. Mol Cell Proteomics 11, 1913–1923. 10.1074/mcp.M112.020586.

Ho, M.Y., Yu, A.L., and Yu, J. (2017). Glycosphingolipid dynamics in human embryonic stem cell and cancer: their characterization and biomedical implications. Glycoconj J 34, 765–777. 10.1007/s10719-016-9715-x.

Jirmo, A.C., Rossdam, C., Grychtol, R., Happle, C., Gerardy-Schahn, R., Buettner, F.F.R., and Hansen, G. (2020). Differential expression patterns of glycosphingolipids and C-type lectin receptors on immune cells in absence of functional regulatory T cells. Immun Inflamm Dis 8, 512–522. 10.1002/iid3.334.

Kannagi, R., Cochran, N.A., Ishigami, F., Hakomori, S., Andrews, P.W., Knowles, B.B., and Solter, D. (1983). Stage-specific embryonic antigens (SSEA-3 and -4) are epitopes of a unique globo-series ganglioside isolated from human teratocarcinoma cells. Embo j 2, 2355–2361. 10.1002/j.1460-2075.1983.tb01746.x.

Kempf, H., Andree, B., and Zweigerdt, R. (2016a). Large-scale production of human pluripotent stem cell derived cardiomyocytes. Adv Drug Deliv Rev 96, 18–30. 10.1016/j.addr.2015.11.016.

Kempf, H., Olmer, R., Haase, A., Franke, A., Bolesani, E., Schwanke, K., Robles-Diaz, D., Coffee, M., Göhring, G., Dräger, G., et al. (2016b). Bulk cell density and Wnt/TGFbeta signalling regulate mesendodermal patterning of human pluripotent stem cells. Nat Commun 7, 13602. 10.1038/ncomms13602.

Konze, S.A., Cajic, S., Oberbeck, A., Hennig, R., Pich, A., Rapp, E., and Buettner, F.F.R. (2017). Quantitative Assessment of Sialo-Glycoproteins and N-Glycans during Cardiomyogenic Differentiation of Human Induced Pluripotent Stem Cells. Chembiochem 18, 1317–1331. 10.1002/cbic.201700100.

Liang, Y.J., Kuo, H.H., Lin, C.H., Chen, Y.Y., Yang, B.C., Cheng, Y.Y., Yu, A.L., Khoo, K.H., and Yu, J. (2010). Switching of the core structures of glycosphingolipids from globo- and lacto- to ganglio-series upon human embryonic stem cell differentiation. Proc Natl Acad Sci U S A 107, 22564–22569. 10.1073/pnas.1007290108.

Liang, Y.J., Yang, B.C., Chen, J.M., Lin, Y.H., Huang, C.L., Cheng, Y.Y., Hsu, C.Y., Khoo, K.H., Shen, C.N., and Yu, J. (2011). Changes in glycosphingolipid composition during differentiation of human embryonic stem cells to ectodermal or endodermal lineages. Stem Cells 29, 1995–2004. 10.1002/stem.750.

Lin, R.J., Kuo, M.W., Yang, B.C., Tsai, H.H., Chen, K., Huang, J.R., Lee, Y.S., Yu, A.L., and Yu, J. (2020). B3GALT5 knockout alters gycosphingolipid profile and facilitates transition to human naïve pluripotency. Proc Natl Acad Sci U S A 117, 27435–27444. 10.1073/pnas.2003155117.

Natunen, S., Satomaa, T., Pitkänen, V., Salo, H., Mikkola, M., Natunen, J., Otonkoski, T., and Valmu, L. (2011). The binding specificity of the marker antibodies Tra-1-60 and Tra-1-81 reveals a novel pluripotency-associated type 1 lactosamine epitope. Glycobiology 21, 1125–1130. 10.1093/glycob/cwq209.

Naujok, O., Kaldrack, J., Taivankhuu, T., Jörns, A., and Lenzen, S. (2010). Selective removal of undifferentiated embryonic stem cells from differentiation cultures through HSV1 thymidine kinase and ganciclovir treatment. Stem Cell Rev Rep 6, 450–461. 10.1007/s12015-010-9148-z.

Ojima, T., Shibata, E., Saito, S., Toyoda, M., Nakajima, H., Yamazaki-Inoue, M., Miyagawa, Y., Kiyokawa, N., Fujimoto, J.I., Sato, T., and Umezawa, A. (2015). Glycolipid dynamics in generation and differentiation of induced pluripotent stem cells. Sci Rep 5, 14988. 10.1038/srep14988.

Okano, H., and Yamanaka, S. (2014). iPS cell technologies: significance and applications to CNS regeneration and disease. Mol Brain 7, 22. 10.1186/1756-6606-7-22.

Pera, M.F., and Trounson, A.O. (2004). Human embryonic stem cells: prospects for development. Development 131, 5515–5525. 10.1242/dev.01451.

Rossdam, C., Konze, S.A., Oberbeck, A., Rapp, E., Gerardy-Schahn, R., von Itzstein, M., and Buettner, F.F.R. (2019). Approach for Profiling of Glycosphingolipid Glycosylation by Multiplexed Capillary Gel Electrophoresis Coupled to Laser-Induced Fluorescence Detection To Identify Cell-Surface Markers of Human Pluripotent Stem Cells and Derived Cardiomyocytes. Anal Chem 91, 6413–6418. 10.1021/acs.analchem.9b01114.

Russo, D., Capolupo, L., Loomba, J.S., Sticco, L., and D’Angelo, G. (2018). Glycosphingolipid metabolism in cell fate specification. J Cell Sci 131. 10.1242/jcs.219204.

Säljö, K., Barone, A., Vizlin-Hodzic, D., Johansson, B.R., Breimer, M.E., Funa, K., and Teneberg, S. (2017). Comparison of the glycosphingolipids of human-induced pluripotent stem cells and human embryonic stem cells. Glycobiology 27, 291–305. 10.1093/glycob/cww125.

Shi, Y., Kirwan, P., and Livesey, F.J. (2012). Directed differentiation of human pluripotent stem cells to cerebral cortex neurons and neural networks. Nat Protoc 7, 1836–1846. 10.1038/nprot.2012.116.

Starzonek, S., Maar, H., Labitzky, V., Wicklein, D., Rossdam, C., Buettner, F.F.R., Wolters-Eisfeld, G., Guengoer, C., Wagener, C., Schumacher, U., and Lange, T. (2020). Systematic analysis of the human tumor cell binding to human vs. murine E- and P-selectin under static vs. dynamic conditions. Glycobiology 30, 695–709. 10.1093/glycob/cwaa019.

Tan, H.L., Fong, W.J., Lee, E.H., Yap, M., and Choo, A. (2009). mAb 84, a cytotoxic antibody that kills undifferentiated human embryonic stem cells via oncosis. Stem Cells 27, 1792–1801. 10.1002/stem.109.

Tang, C., Lee, A.S., Volkmer, J.P., Sahoo, D., Nag, D., Mosley, A.R., Inlay, M.A., Ardehali, R., Chavez, S.L., Pera, R.R., et al. (2011). An antibody against SSEA-5 glycan on human pluripotent stem cells enables removal of teratoma-forming cells. Nat Biotechnol 29, 829–834. 10.1038/nbt.1947.

Tateno, H., Toyota, M., Saito, S., Onuma, Y., Ito, Y., Hiemori, K., Fukumura, M., Matsushima, A., Nakanishi, M., Ohnuma, K., et al. (2011). Glycome diagnosis of human induced pluripotent stem cells using lectin microarray. J Biol Chem 286, 20345–20353. 10.1074/jbc.M111.231274.

Thiesler, C.T., Cajic, S., Hoffmann, D., Thiel, C., van Diepen, L., Hennig, R., Sgodda, M., Weiβmann, R., Reichl, U., Steinemann, D., et al. (2016). Glycomic Characterization of Induced Pluripotent Stem Cells Derived from a Patient Suffering from Phosphomannomutase 2 Congenital Disorder of Glycosylation (PMM2-CDG). Mol Cell Proteomics 15, 1435–1452. 10.1074/mcp.M115.054122.

Wright, A.J., and Andrews, P.W. (2009). Surface marker antigens in the characterization of human embryonic stem cells. Stem Cell Res 3, 3–11. 10.1016/j.scr.2009.04.001.

Zhou, D., Henion, T.R., Jungalwala, F.B., Berger, E.G., and Hennet, T. (2000). The beta 1,3-galactosyltransferase beta 3GalT-V is a stage-specific embryonic antigen-3 (SSEA-3) synthase. J Biol Chem 275, 22631–22634. 10.1074/jbc.C000263200.

